# Memory Replay in Balanced Recurrent Networks

**DOI:** 10.1101/069641

**Authors:** Nikolay Chenkov, Henning Sprekeler, Richard Kempter

**Affiliations:** Institute for Theoretical Biology, Humboldt-Universität zu Berlin, Berlin, Germany; Department of Software Engineering and Theoretical Computer Science, Technische Universität Berlin, Berlin, Germany; Bernstein Center for Computational Neuroscience Berlin, Berlin, Germany

## Abstract

Complex patterns of neural activity appear during up-states in the neocortex and sharp waves in the hippocampus, including sequences that resemble those during prior behavioral experience. The mechanisms underlying this replay are not well understood. How can small synaptic footprints engraved by experience control large-scale network activity during memory retrieval and consolidation? We hypothesize that sparse and weak synaptic connectivity between Hebbian assemblies are boosted by pre-existing recurrent connectivity within them. To investigate this idea, we connect sequences of assemblies in randomly connected spiking neuronal networks with a balance of excitation and inhibition. Simulations and analytical calculations show that recurrent connections within assemblies allow for a fast amplification of signals that indeed reduces the required number of inter-assembly connections. Replay can be evoked by small sensory-like cues or emerge spontaneously by activity fluctuations. Global—potentially neuromodulatory—alterations of neuronal excitability can switch between network states that favor retrieval and consolidation.

**Author Summary:** Synaptic plasticity is the basis for learning and memory, and many experiments indicate that memories are imprinted in synaptic connections. However, basic mechanisms of how such memories are retrieved and consolidated remain unclear. In particular, how can one-shot learning of a sequence of events achieve a sufficiently strong synaptic footprint to retrieve or replay this sequence? Using both numerical simulations of spiking neural networks and an analytic approach, we provide a biologically plausible model for understanding how minute synaptic changes in a recurrent network can nevertheless be retrieved by small cues or even manifest themselves as activity patterns that emerge spontaneously. We show how the retrieval of exceedingly small changes in the connections across assemblies is robustly facilitated by recurrent connectivity within assemblies. This interaction between recurrent amplification within an assembly and the feed-forward propagation of activity across the network establishes a basis for the retrieval of memories.

## Introduction

The idea of sequential activation of mental concepts and neural populations has deep roots in the history of the cognitive sciences [12, 88, 97] as well as its share of criticism [56]. In one of the most influential works in neuroscience, Donald Hebb extended this concept by suggesting that neurons that fire simultaneously should be connected to each other, thus forming a cell assembly that represents an abstract mental concept [41]. He also suggested that such assemblies could be connected amongst each other, forming a network of associations in which one mental concept can ignite associated concepts by activating the corresponding assemblies. Hebb referred to the resulting sequential activation as well as the underlying circuitry as “phase sequence”. We will refer to such connectivity patterns as “assembly sequences”.

The notion of Hebbian assemblies has triggered a huge number of experimental studies (reviewed in [96]), but relatively few experiments have been dedicated to the idea of assembly sequences [2, 53]. Many theoretical studies focused on feedforward networks, also known as synfire chains [1, 7, 25, 44]. Synfire chains are characterized by a convergent-divergent feedforward connectivity between groups of neurons, where pulse packets of synchronous firing can propagate through the network. Few works were also dedicated on synfire chains embedded in recurrent networks [6, 54, 89], however, without explicitly considering recurrent connectivity within groups.

In this study, we combine the concept of feedforward synfire chains with the notion of recurrently connected Hebbian assemblies to form an assembly sequence. Using numerical simulations of spiking neural networks, we form assemblies consisting of recurrently connected excitatory and inhibitory neurons. The networks are tuned to operate in a balanced regime where large fluctuations of the mean excitatory and inhibitory input currents cancel each other. In this case, distinct assemblies that are sparsely connected in a feedforward fashion can reliably propagate transient activity. This replay can be triggered by external cues for sparse connectivities, but also can be evoked by background activity fluctuations for larger connectivities. Modulating the population excitability can shift the network state between cued-replay and spontaneous-replay regimes. Such spontaneous events may be the basis of the reverberating activity observed in the neocortex [18, 49, 63] or in the hippocampus [26, 58, 86]. Finally, we show that assembly sequences can also be replayed in a reversed direction (i.e., reverse replay) as observed during replay of behavior sequences [23, 30].

## Results

To test Hebb’s hypothesis on activity propagation within a recurrent network, we use a network model of excitatory and inhibitory conductance-based integrate-and-fire neurons. The network has a sparse random background connectivity *p*_rand_ = 0.01. We form a neural assembly (Fig 1A) by picking *M* excitatory ( *M* = 500 if not stated otherwise) and *M*/4 inhibitory neurons and connecting them randomly with probability *p*_rc_, resulting in a mutually coupled excitatory and inhibitory population. The new connections are created independently and in addition to the background connections. To embed an assembly sequence in the network, we first form 10 non-overlapping assemblies. The assemblies are then connected in a feedforward manner where an excitatory neuron from one group projects to an excitatory neuron in the subsequent group with probability *p*_ff_ (Fig 1B). Thus, by varying the feedforward and the recurrent connectivities, we can set the network structure anywhere in the spectrum between the limiting cases of synfire chains (*p*_ff_ > 0, *p*_rc_ = 0) and uncoupled Hebbian assemblies (*p*_ff_ = 0, *p*_rc_ > 0), as depicted in Fig 1C.

**Fig 1.**
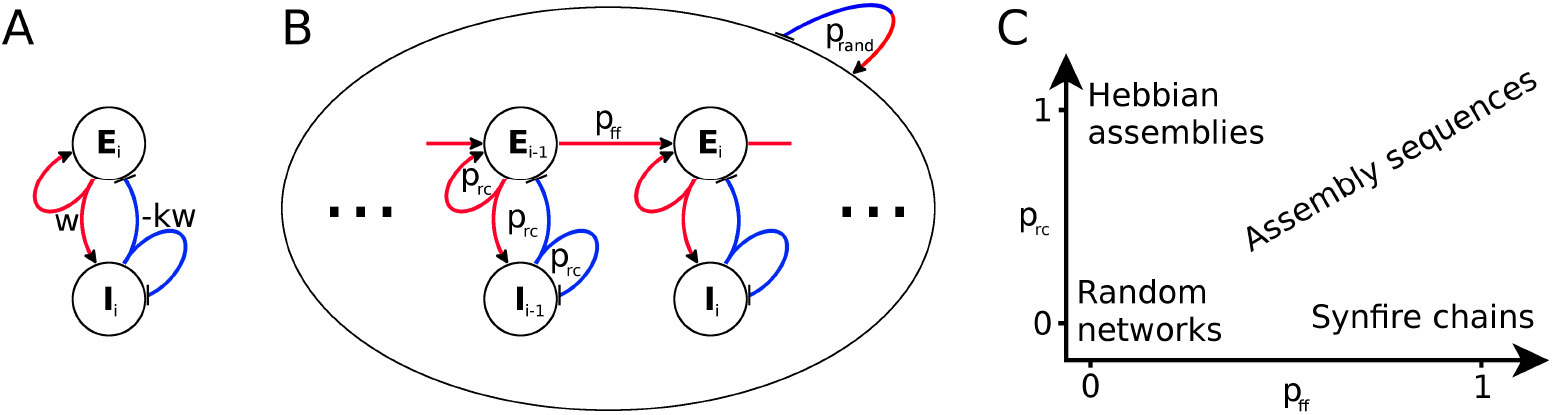
Network connectivity. **A:**Schematic of an assembly *i* consisting of an excitatory (*E_i_*) and an inhibitory (*I_i_*) population. Red and blue lines indicate excitatory and inhibitory connections, respectively. The symbols w and −kw denote total synaptic couplings between populations. **B:** Sketch of network connectivity. The inhomogeneous network is randomly connected with connection probability *p_rand_*. A feedforward structure consisting of 10 assemblies (only *i* − 1 and *i* shown) is embedded into the network. Each assembly is formed by recurrently connecting its neurons with probability *p*_rc_. Subsequent assemblies are connected with feedforward probability *p*_ff_ between their excitatory neurons. **C:** Embedded structure as a function of connectivities.

To ensure that the spontaneous activity of the network is close to an *in-vivo* condition, we use Hebbian plasticity of inhibitory connections [94], which has been shown to generate a balance of excitatory and inhibitory currents in individual neurons (Fig 2A). As a consequence, spikes are caused by current fluctuations (Fig 2B), and the network settles into a state of asynchronous irregular (AI) firing (Fig 2C, 0-500 ms). If we then stimulate the first group in the embedded assembly sequence (Fig 2C, 500 ms), the network responds with a wave of activity that traverses the whole assembly sequence, as hypothesised by Hebb [41]. We refer to such a propagation of activity wave as replay. As excitatory and inhibitory neurons are part of the assemblies, they both have elevated firing rates during group activation ( ∼ 100 spikes/sec for excitatory, and ∼ 60 spikes/sec for inhibitory neurons). Because excitatory neurons in the assembly sequence transiently increase their firing rates from 5 to 100 spikes/sec, a replay can be inferred from their large change in activity, which resembles replay in hippocampal CA networks [58]. On the other hand, interneurons have higher background firing rates of ∼ 20 spikes/sec and smaller maximum firing rates of ∼ 60 spikes/ sec during replay. As a result, interneurons have a much lower ratio of peak to background activity than excitatory neurons in our model, in line with lower selectivity of interneurons (e.g., [100]).

**Fig 2.**
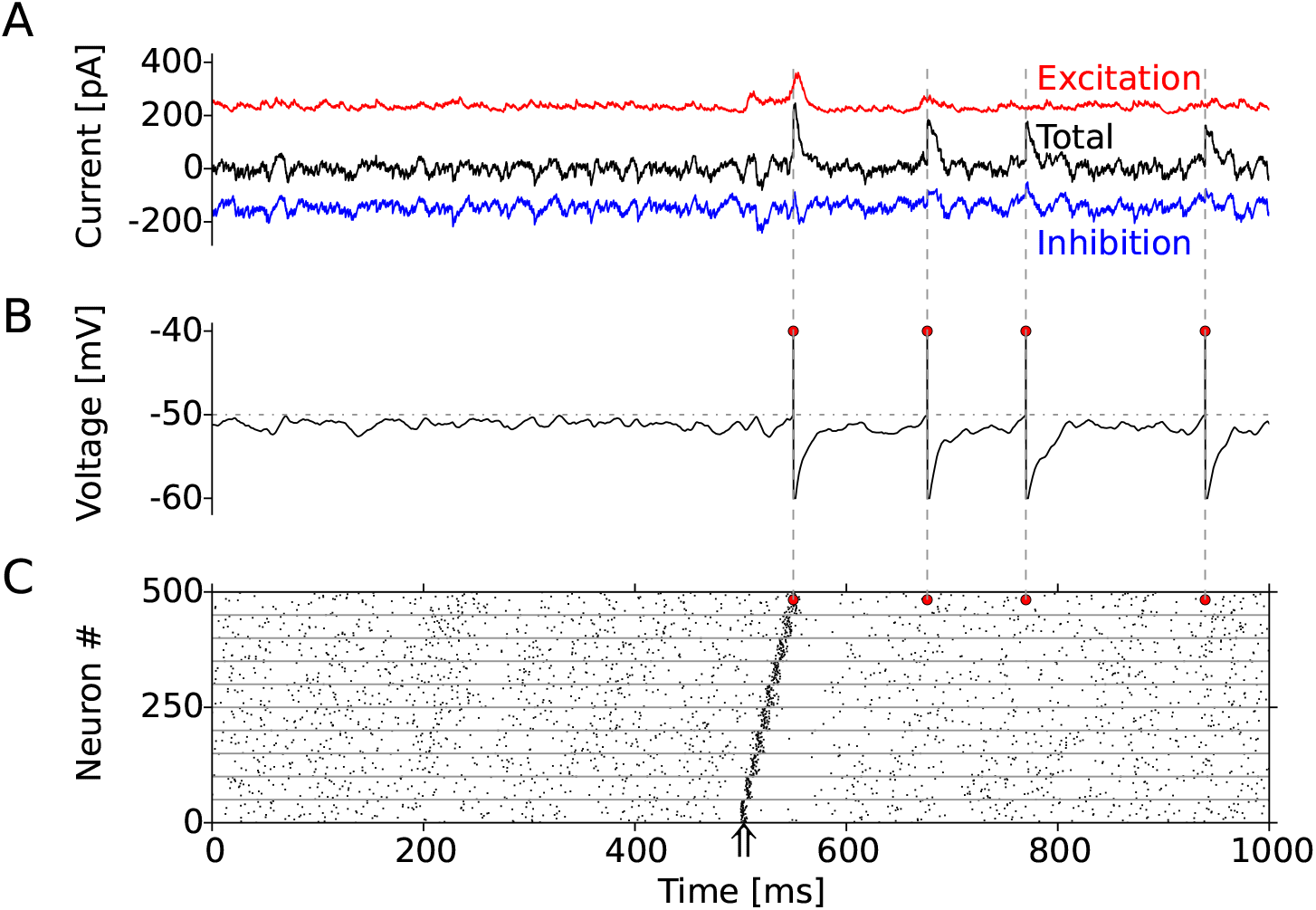
Example of activity in a balanced network. **A:** Input currents experienced by an example neuron. The excitatory input is denoted by the red trace while the inhibitory one is in blue. The black curve shows the sum of all currents: synaptic, injected, and leak currents. **B:** Membrane potential of the same neuron. Red dots denote the times of firing. **C:** Raster plot of spikes times of 500 neurons for 1 second. For better readability, only 50 neurons (out of 500) per group are shown. At time 500 ms, the first group is stimulated (arrow) and the activity propagates through the assembly sequence, resulting in a replay. The red dots correspond to the firing of the example neuron in A and B.

### Sparse feedforward connectivity is sufficient for replay

Whether an assembly sequence is replayed is largely determined by the connectivities within and between assemblies. Therefore, we first study how the quality of replay depends on the recurrent (*p*_rc_) and the feedforward (*p*_ff_) connectivities. The network dynamics can be roughly assigned to regimes where the connectivity is too weak, strong enough, or too strong for a successful replay. We use a quality measure of replay (for details see Materials and Methods), which determines whether activations of the first group propagate to the end of the sequence without evoking a “pathological” burst of activity (Fig 3).

**Fig 3.**
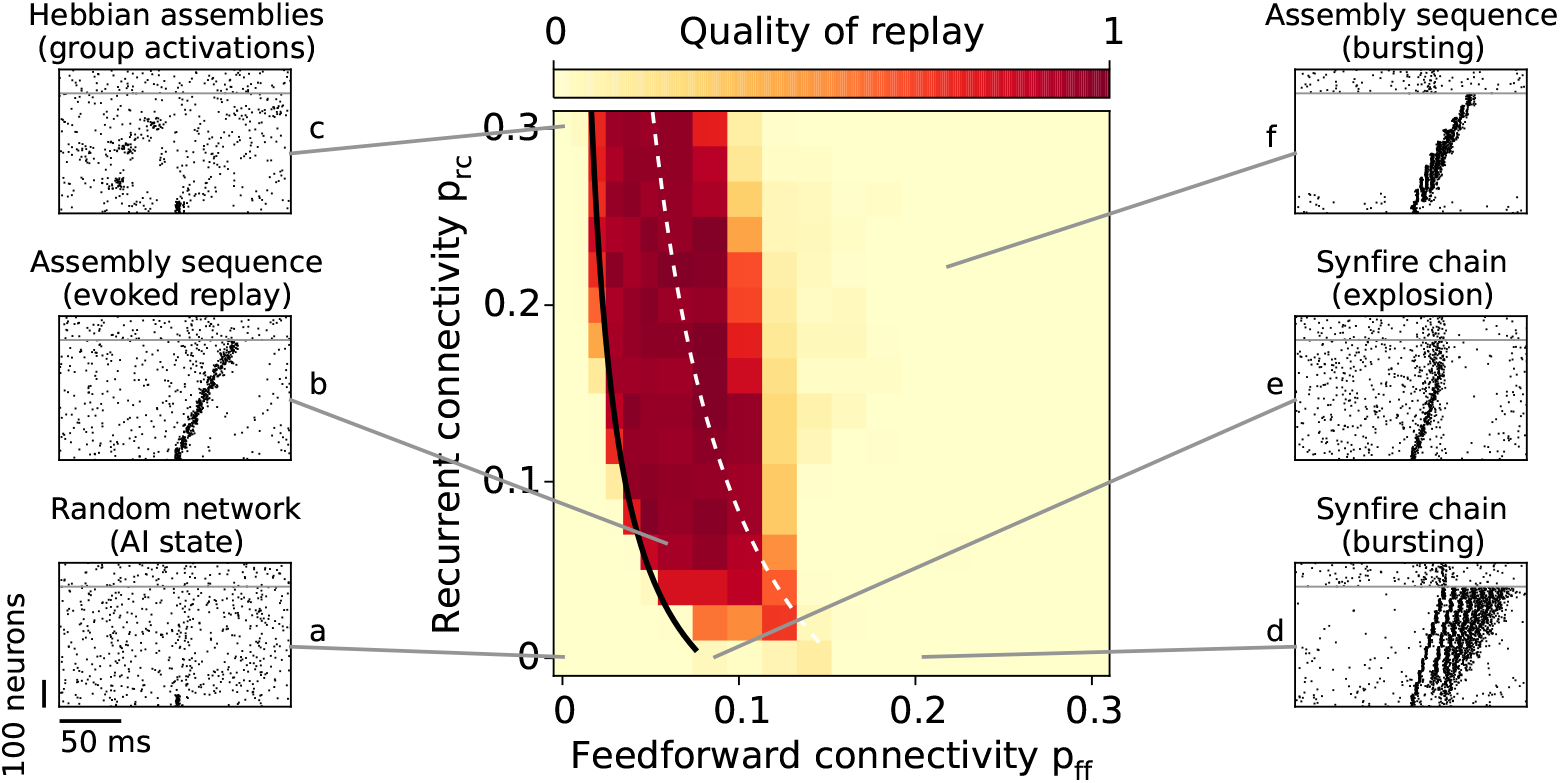
Evoked replay. Assembly-sequence activation as a function of the feedforward *p* ff and the recurrent *p*_rc_ connectivities. The color code denotes the quality of replay, that is, the number of subsequent groups firing without bursting (see Materials and Methods). The black curve corresponds to the critical connectivity required for a replay where the slope *c* of the transfer function (See Materials and Methods and Eq 1) is matched manually to fit the simulation results for connectivities *p*_rc_ = 0.08 and *p*_ff_ = 0.04. The slope *c* is also estimated analytically (dashed white line). The raster plots (**a-f**) illustrate the dynamic regimes observed for different connectivity values;neurons above the gray line belong to the background neurons.

Naturally, for a random network (*p*_ff_ = 0, *p*_rc_ = 0, Fig 3a) the replay fails because the random connections are not sufficient to drive the succeeding groups. In the case of uncoupled Hebbian assemblies (e.g., *p*_ff_ = 0, *p*_rc_ = 0.30), groups of neurons get activated spontaneously (Fig 3c), which is reminiscent to the previously reported cluster activation [61] but on a faster time scale. Already for sparse connectivity (e.g., *p*_ff_ = *p*_rc_ = 0.06) the assembly-sequence replay is successful (Fig 3b). In the case of denser recurrence (*p*_rc_ ≈ 0.10), a pulse packet propagates for even lower feedforward connectivity (*p*_ff_ ≈ 0.03). The feedforward connectivity that is required for a successful propagation decreases with increasing recurrent connectivity because assemblies of excitatory and inhibitory neurons can increase small fluctuations of the input through “balanced amplification” [42, 72].

For high feedforward 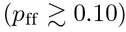 but low recurrent 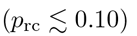 connectivity, the replay has low quality. In this case, excitatory neurons receive small recurrent inhibitory input compared to the large feedforward excitation, because the recurrent connection probability is lower than the feedforward one. Due to this lack of sufficient inhibitory input, the propagating activity either leads to run-away excitation (Fig 3e), also called synfire explosion [5, 66], or to epileptiform bursting (Fig 3d). When both recurrent and feedforward connectivities are high, the inhibition is able to keep the propagating activity transient (Fig 3f). However, due to the strong input each neuron is firing multiple times within a small time window. Due to this bursting, the replay has a low quality.

To get an analytical understanding of the network, we use a linear approximation of the network dynamics to derive conditions under which replay is successful. The key determinant for replay is an amplification factor 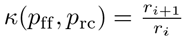, which measures how large is the rate *r_i+1_* in group *i*+ 1 in relation to the rate in the previous group *i*.

In the case where the amplification factor is smaller than one (*r_i+1_* < *r_i_*), the activity propagating through the assembly sequence will decrease at each step and eventually vanish, while for amplification larger than one (*r_i+1_* > *r_i_*) one would expect propagating activity that increases at each step in the sequence. An amplification factor *κ*(*p*_ff_, *p*_rc_) = 1 represents the critical value of connectivity for which the replay is stable, and the magnitude of activations is similar across groups. In the Materials and Methods we show that a linear model can approximate the amplification factor by

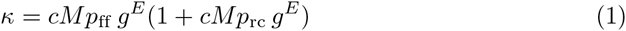

where *c* = 0.25 nS^−1^ is a constant that fits the model to the data (see Materials and Methods). We can interpret *к* as an “effective feedforward connectivity” because the recurrent connectivity (*p*_rc_) effectively scales up the feedforward connectivity *p*_ff_. We can match the analytical results for critical connectivities to the numerical simulation, and show a qualitative fit between the approaches (black line in Fig 3).

We note that the number of excitatory synapses that is needed for an association, *M*^2^(*p*_rc_ + *p*_ff_), weakly depends on the position on the line *к* = 1. By solving argmin *p*_rc_, *p*_ff_ ∈к = 1 *M*^2^(*p*_rc_ + *p*_ff_) we find that the minimum number of new connections required for a replay is obtained for *p*_rc_ = 0 because lines for which *p*_rc_ + *p*_ff_ = const have slope of −1 in Fig 3, and the slope of the line defined by *к* = 1 has a more negative slope. For example, when *p*_rc_ = 0.0, we need 40 new synapses; for *p*_rc_ = 0.05, we need 50 new synapses; and for *p*_rc_ = 0.2, 111 synapses are required for a new association. However, as feedforward connections might be created/facilitated on demand in one-shot learning, it is advantageous to keep their number low at the cost of higher recurrent connectivity, which has more time to develop *prior* to the learning. We extend this arguments further in the Discussion.

In summary, the recurrent connections within an assembly play a crucial role in integrating and amplifying the input to the assembly. This facilitation of replay is predominantly due to the excitatory-to-excitatory (E-E) recurrent connections, and not due to the excitatory-to-inhibitory (E-I) connections, a connectivity also known as “shadow pools” [5]. Embedding shadow pools and omitting the E-E connectivity within assemblies has no beneficial effect on the quality of replay (results not shown).

### Recurrent connections are important for pattern completion

Neural systems have to deal with obscure or incomplete sensory cues. A widely adopted solution is pattern completion, that is, reconstruction of patterns from partial input. We examine how the network activity evolves in time for a partial or asynchronous activation of the first assembly.

To determine the capability of our network to complete patterns, we quantify the replay when only a fraction of the neurons in the first group is stimulated by external input. If 60 % of the neurons in the first group (strong cue) are synchronously activated (Fig 4A, left panel), the quality of replay is virtually the same as in the case of full stimulation (100% activated) in Fig 3. However, when only 20 % of the neurons (weak cue) are simultaneously activated (Fig 4A, middle panel), we see a deterioration of replay mostly for low recurrent connectivities. The effect of the recurrent connections is illustrated in the right-most panel in Fig 4A where quality of replay is shown as a function of *p*_rc_ while the feedforward connectivity was kept constant (*p*_ff_ = 0.05).

**Fig 4.**
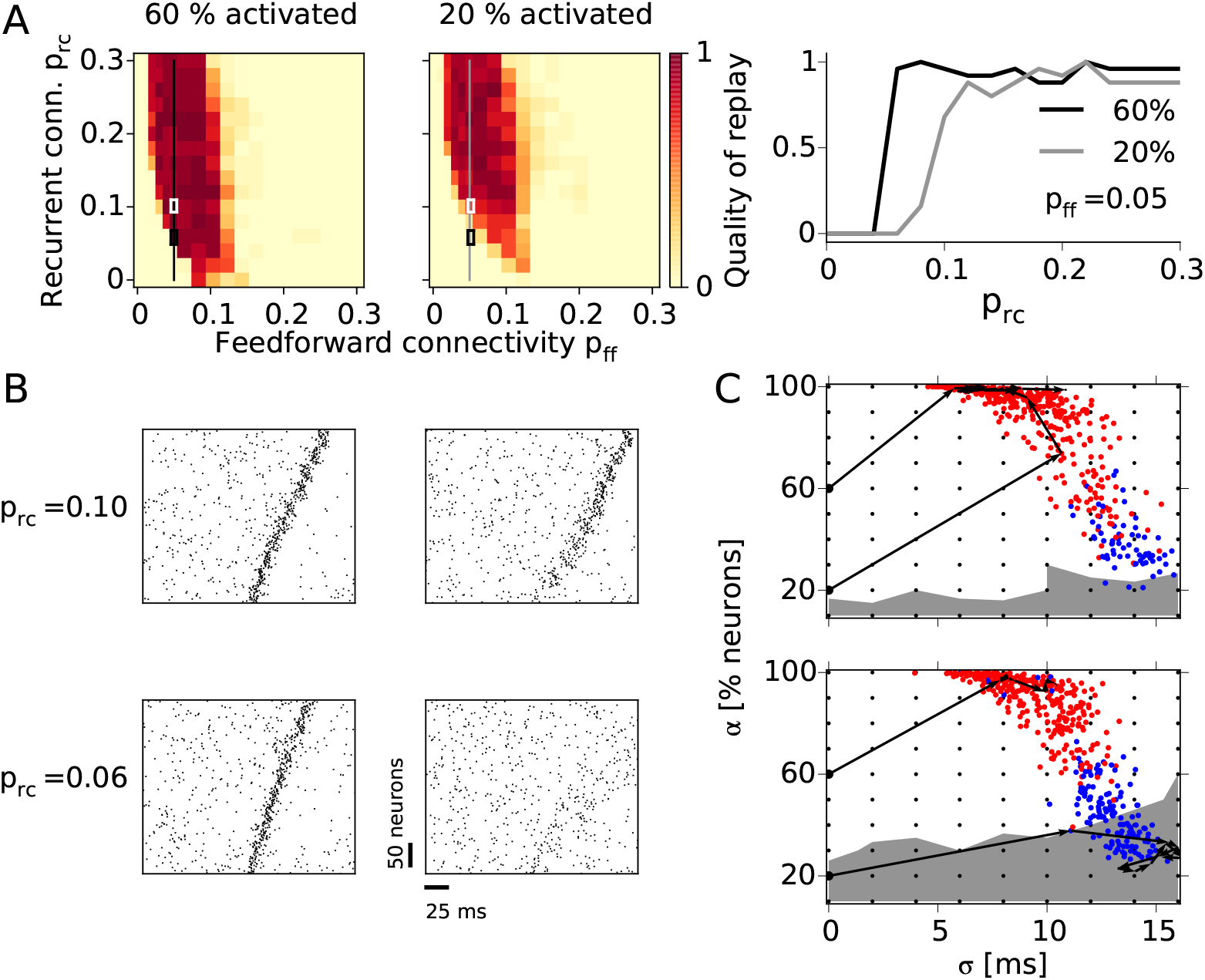
Pattern completion. **A:** Quality of replay after partial activation of the first group for cue size 60% (left panel) and 20% (middle) as a function of feedforward and recurrent connectivity. The right-most panel shows the quality replay after a cue activation (20% and 60%) as a function of the recurrent connectivity (*p*_rc_) while the feedforward connectivity is constant (*p*_ff_ = 0.05). **B:** Examples of network activity during 60% (left) and 20% (right) cue activation. The top and bottom raster plots correspond to assembly sequences with higher (*p*_rc_ = 0.10, top) and lower (*p*_rc_ = 0.06, bottom) recurrent connectivity, highlighted in A with white and black rectangles, respectively. **C:** State-space portraits representing the pulse-packet propagation. The activity in each group is quantified by the fraction of firing excitatory neurons (*α*) and the standard deviation of their spike times (*σ*). The initial stimulations are denoted with small black dots while the colored dots denote the response of the first group to the stimulations; red dot if the whole sequence is activated, and blue otherwise. Stimulations in the region with white background result in replays, while stimulating in the gray region results in no replay. The black arrows illustrate the evolution of pulse packets during the replays in B. Top: *p*_rc_ = 0.10; bottom: *p*_rc_ = 0.06.

Small input cues lead to a weak activation of the corresponding assembly. In the case of stronger connectivity (e.g., *p*_rc_) this weak activity can build up and result in a replay as shown in the example from Fig 4B. The top and bottom rows of raster plots correspond to two assembly sequences with different recurrent connectivities, as highlighted by the rectangles in Fig 4A, while left and right columns show the activity during strong and weak cues, respectively. In the case of *p*_ff_ = 0.05 and *p*_rc_ = 0.10 (Fig 4B, top-right), the weak cue triggers a wide pulse packet with large temporal jitter in the first groups, which gradually shapes into a synchronous pulse packet as it propagates through the network. On the other hand, for a smaller recurrent connectivity (*p*_rc_ = 0.06), the 20% partial activation triggers a rather weak response that does not result in replay (Fig 4B, bottom-right).

The quality of replay depends not only on the number of neurons that are activated but also on the temporal dispersion of the pulse packet. Here, we adopt a quantification method that represents the activity evolution in a state-space portrait [25]. Fig 4C shows the time course of the fraction *α* of cells that participate in the pulse packet and the temporal dispersion *σ* of the packet as the pulse propagates through the network. The state-space representation of two assembly sequences with equal feedforward (*p*_ff_ = 0.05) but different recurrent connectivity are shown in Fig 4C (top: *p*_rc_ = 0.10, bottom: *p*_rc_ = 0.06). For each assembly sequence we repeatedly stimulated the first group with varying cue size *α* and time dispersion *σ*, depicted by the black dots. Depending on the strength and dispersion of the initial stimulation, the dynamics of a network can enter one of two attractor points. For high *α* and low *σ* the pulse packet propagates, entering the so-called synfire attractor (white background). On the other hand, for low *α* and high *σ* the pulse packet dies out resulting in low asynchronous firing (gray background). The black-arrow traces in Fig 4C are example trajectories that describe the propagating pulse packets from Fig 4B in the (*α − σ*) space.

To summarize, increasing both the recurrent and feedforward connectivity facilitates the replay triggered by weak and dispersed inputs. Recurrent connectivity is particularly important for pattern completion.

### Spontaneous replay

An interesting feature of assembly sequences is the potential emergence of spontaneous activations, that is, a replay when no specific input is given to the network. Random fluctuations in the network can be amplified by the feedforward structure and give rise to a spontaneous wave of propagation.

We find that spontaneous and evoked replay share various features such as sequential group activation on the background of AI network activity (Fig 5A, rasters a and b). As in the case of evoked replay, for exceedingly large connectivities the network dynamics can be dominated by epileptiform bursting activity (Fig 5A, rasters c and d).

**Fig 5.**
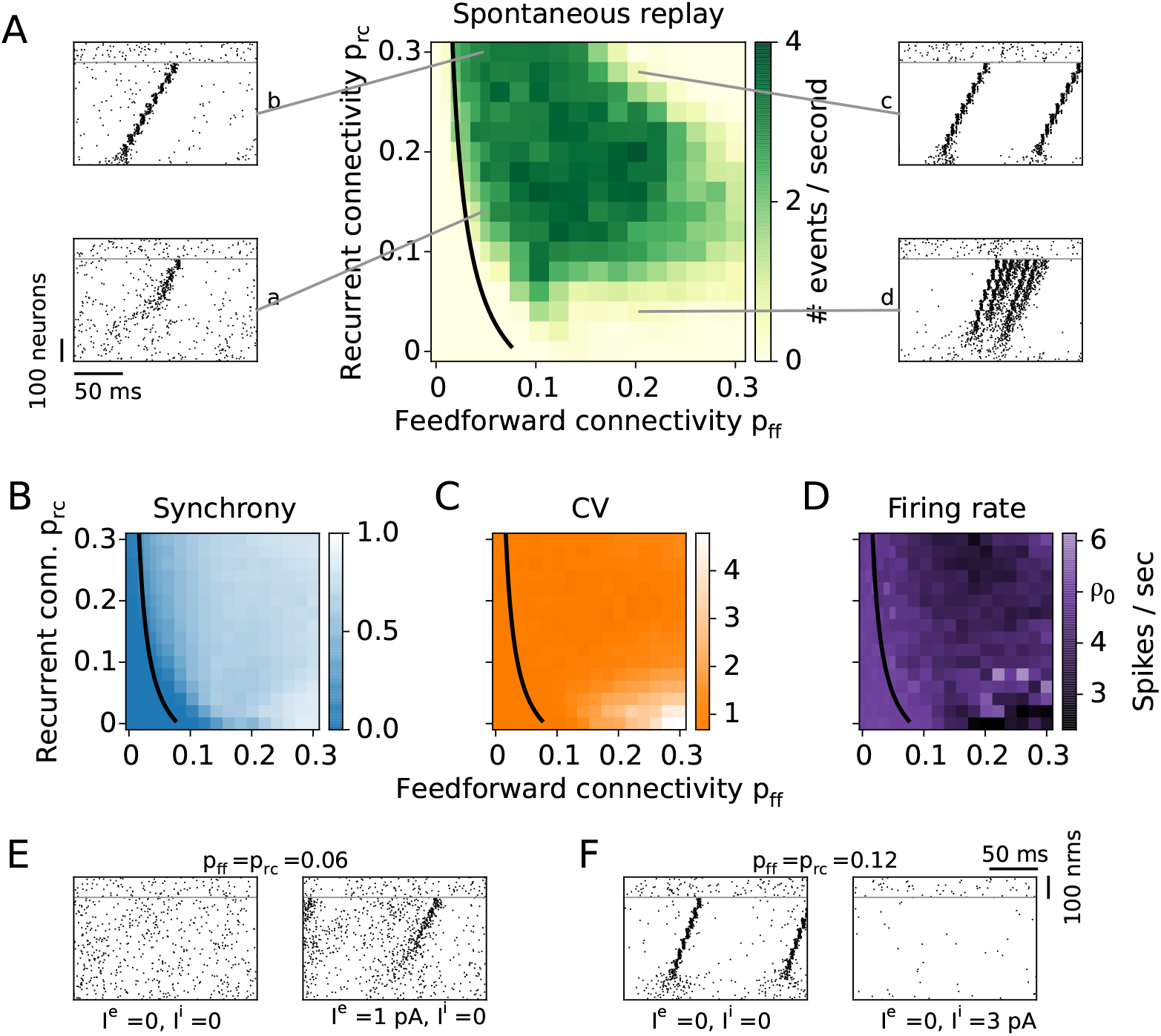
Spontaneous network activity. **A:** The rate of spontaneous sequence activation is measured in the unperturbed network. The black curve is the analytical result for the lower bound of successful propagation from Fig 3. Examples of spontaneous replays for different connectivities are shown in the raster plots **a-d**. Synchrony (**B**), coefficient of variation (**C**), and firing rate (**D**) are averaged over the neurons in the last group of the sequence. **E:** Spontaneous events modulated by an external input. For low enough connectivities no spontaneous events occur (left). A small additional constant current input to the whole excitatory population ( *I^e^* = 1 pA) generates spontaneous replays (right). **F:** A densely connected network shows replays (left). Once the inhibitory population receives an additional constant current input ( *I^i^* = 3 pA), the firing rate decreases and no spontaneous events occur (right).

To assess spontaneous replay, we quantify the number of replay events per time taking into account their quality, i.e., huge bursts of propagating activity are disregarded as replay. The rate of spontaneous activation increases as a function of both the feedforward (*p*_ff_) and the recurrent (*p*_rc_) connectivity (Fig 5A). For large connectivities (*p*_ff_, *p*_rc_ > 0.20) the quality of the spontaneous events is again poor and mostly dominated by strong bursts (Fig 5A, raster c). The dynamics of networks with large feedforward and low recurrent connections is dominated by long-lasting bursts of activity consisting of multiple sequence replays within each burst (Fig 5A, raster d). The maximum rate of activations does not exceed 4 events per second because the inhibitory synaptic plasticity adjusts the inhibition such that the excitatory firing rate is close to 5 spikes/sec.

To better characterize spontaneous dynamics, we refer to more extensive measures of the network dynamics. First, to account for deviations from the AI network state, we measure the synchrony of firing among neurons within the assemblies. To this end, we calculate the average pairwise correlation coefficient of spike trains of neurons within the same group. A low synchrony (value ∼ 0) means that neurons are uncorrelated, while a high synchrony (value ∼ 1) reveals that neurons fire preferentially together and seldom (or not at all) outside of an assembly activation. Because the synchrony builds up while activity propagates from one group to the next, a synchronization is most pronounced in the latter groups of the sequence. Therefore, we use correlations within the last group of the sequence as a measure of network synchrony (Fig 5B). The average synchrony is low ( ∼ 0) for low connectivities (*p*_ff_, *p*_rc_ < 0.10) and increases as a function of both p_ff_ and *p*_rc_. In the case of high *p*_rc_, neurons participating in one assembly excite each other, and hence tend to fire together. On the other hand, for high *p*_ff_, neurons within an assembly receive very similar input from the preceding group, so they fire together. This attachment of single neurons to group activity has two major consequences: first, it alters the AI state of the network, and second, it alters the stochastic behavior of the neurons, leading to more deterministic firing and bursting.

The network exhibits frequent epileptiform bursting in the case of high feedforward and low recurrent connectivities (raster plot examples in Fig 3, panel d, and Fig 5A, panel d). To assess this tendency of neurons to fire in bursts, we calculate the coefficient of variation (CV) for individual neurons’ spike trains. The average CV of neurons in the last group of the sequence exhibits Poisson-like irregular firing (CV value ∼ 1) for a large range of parameters (Fig 5C). However, for high *p*_ff_ ( ≥0.10) and low *p*_rc_ ( ≤0.10), the CV value exceeds 1, in line with irregular and bursting firing. In this parameter region, small fluctuations of activity in the first groups of the sequence are strongly amplified by the underlying feedforward connectivity, leading to ever increasing activity in the following groups (Fig 5A, panel d). Because of the variable shapes and sizes of these bursts, they are not always classified as spontaneous activations in Fig 5A. Highly bursty firing (CV > 3) and high synchrony ( ∼ 1) suggest that the network cannot be properly balanced.

To test whether the inhibitory plasticity can balance the network activity when assembly sequences are embedded, we measure the average firing rate in the last group of the sequence (Fig 5D). The firing rate deviates from the target rate of 5 spikes/sec mostly for high feedforward connectivity 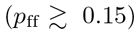. This inability of inhibition to keep the firing rate at the target value can be explained by the frequent replays that shape a stronger inhibitory input during the balancing of the network. Once the inhibition gets too strong, neurons can fire only when they receive excessive amount of excitation. Thus, in the case of high clustering, e.g., strong assembly connectivity, the inhibitory plasticity prevents the neurons from reaching high firing rates, but is unable to sustain an AI state of the network.

### Control of spontaneous and cued replay by external input

Further, we investigate how spontaneous and cued replay are related. The black line in Fig 5A refers to the analytical approximation for connectivities that enable evoked replay. Compared to the connectivity region of successfully evoked replays in Fig 3, the region for spontaneous replays in Fig 5 is slightly shifted to the top and to the right. Therefore, in only a narrow area of the parameter space, sequences can be replayed by external input but do not get spontaneously activated. This finding suggests that to embed a sequence with high signal-to-noise ratio of propagation, the connectivities should be chosen appropriately, in line with previous reports [55]. In what follows we show that the size of this region can be controlled by external input to the network.

Fig 5E and F illustrate how a small amount of global input current to all excitatory or all inhibitory neurons can modulate the network and shift it between AI and spontaneous-replay regimes. In the first example, the connectivities are relatively low (*p*_ff_ = *p*_rc_ = 0.06) such that replay can be evoked (Fig 3) but no spontaneous activations are present (Fig 5A and Fig 5E, left). After injecting a small additional current of only 1 pA into the whole excitatory population, the network becomes more excitable, i.e., the firing rate rises from 5 to 12 spikes/ sec and spontaneous replays do arise (Fig 5E, right).

On the other hand, in a network with high connectivities (*p*_ff_ = *p*_rc_ = 0.12), replay can be reliably evoked (Figs 3 and 4A) and also occurs spontaneously (Fig 5A). An additional input current of 3 pA to the inhibitory population decreases the firing rate of the excitatory population from 5 to 0.33 spikes/sec and shifts the network from a regime showing frequent spontaneous replays to a no-replay, AI regime (Fig 5F, left and right, respectively). Nevertheless, replays can still be evoked as in Fig 3 (result not shown). Hence, the spontaneous-replay regime and the average firing rate in the AI state can be controlled by global or unspecific external current.

In summary, the balanced AI network state and successfully evoked replay of assembly sequences can coexist for a range of connectivities. For higher connectivities, the underlying network structure amplifies random fluctuations, leading to spontaneous propagations of activity between assemblies. A dynamical control of the rate of spontaneous events is possible through external input, which modulates the network activity and excitability. In the brain, such a switching between regimes could be achieved via neuromodulators, in particular via the cholinergic or adrenergic systems [40, 87].

### Smaller assemblies require higher connectivity

So far, we have shown basic properties of sequences at fixed assembly size *M* = 500. To determine the role of this group size in replay, we vary *M* and the connectivity while keeping the size of the network fixed. As we have already explored how recurrent and feedforward connections determine replay individually, we now consider the case where they are equal, i.e., *p*ff = *p*_rc_ = *p*.

Assembly sequences can be successfully replayed after stimulation for various assembly sizes (Fig 6A). Smaller assemblies require denser connectivity (e.g., *p* = 0.25 for *M* = 100), while larger assemblies allow sparser connectivity (e.g., *p* = 0.05 for *M* = 500). Moreover, assemblies as small as 20 neurons are sufficient to organize a sequence given the condition of all-to-all connectivity within and between assemblies (result not shown). The analytically derived critical value of effective connectivity *к* = 1 is in agreement with the numerical simulations (black line in Fig 6A).

**Fig 6.**
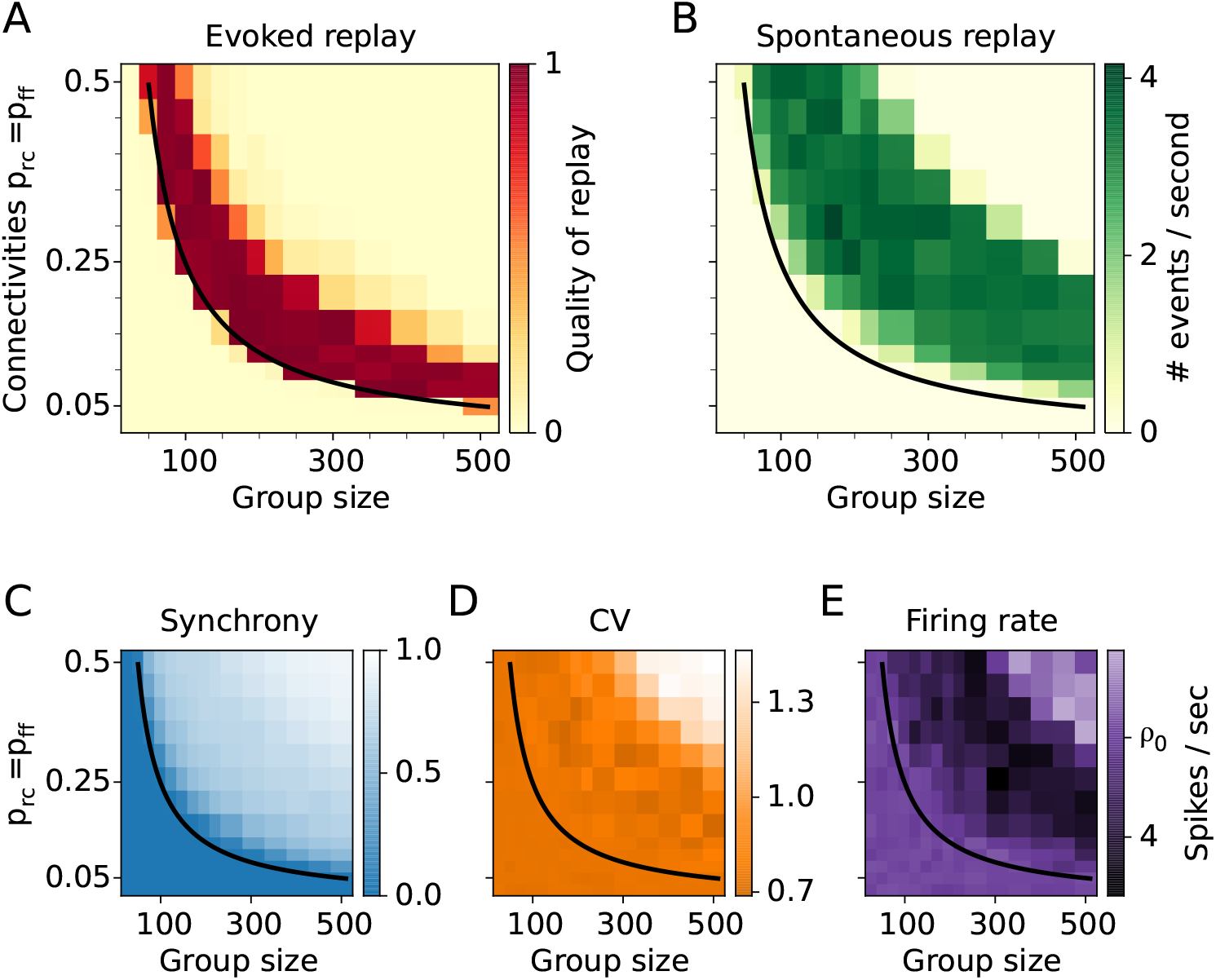
Assembly-sequence activation for various group sizes and connectivities. **A:**Simulation results for the quality of replay. **B:** Rate of spontaneous replay. **C:** Synchrony. **D:** Coefficient of variation **E:** Firing rate. *ρ*_0_ = 5 spikes/sec is the target firing rate. In C, D, and E quantities are averaged over the neurons in the last group of the sequence. The black line is an analytical estimate for the evoked replay as in Figs 3 and 5.

To further characterize the network dynamics for varying group size, we measure the rate of spontaneous activations of assembly sequences in undisturbed networks driven solely by constant input. Fig 6B indicates that spontaneous replays occur for a limited set of parameters resembling a banana-shaped region in the (*M*, *p*) plane. The parameter region for spontaneous replays partly overlaps with that of evoked replay. Again, there is a narrow range of parameters to the right of the black line in Fig 6B for which sequences can be evoked by external input while not being replayed spontaneously. As shown above, the size of this region can be controlled by external input to the whole network (Fig 5E,F).

To further assess the spontaneous dynamics, we measure the firing synchrony of neurons within the last group. The synchrony grows as function of both connectivity and group size (Fig 6C). The fact that the synchrony approaches the value one for higher connectivity and group size indicates that the network dynamics gets dominated by spontaneous reactivations. The simulation results reveal that neurons always fire rather irregular with CVs between 0.7 and 1.4 (Fig 6D). Because the recurrent and the feedforward connectivities are equal (*p*_ff_ = *p*_rc_ = *p*), the inhibition is always strong enough and does not allow epileptiform bursting activity. This behavior is reflected in a rather low maximal value of the coefficient of variation (CV<1.4) compared to the results from Fig 5, where the CV could exceed values of 4 for low *p*_rc_. The measured firing rates in the last assembly are at the target firing rate of *ρ*_0_ = 5 spikes/sec for parameter values around and below the critical value *к* = 1 (Fig 6E). However, for increasing connectivity *p* and increasing group size *M*, the firing rate deviates from the target, indicating that the inhibitory plasticity cannot keep the network fully balanced.

To conclude, the assembly size *M* plays an important role in the network activity. The critical values of connectivity and group size for successful propagation are inversely proportional. Thus, the analytics predicts that larger assemblies of several thousands neurons require only a fraction of a percent connectivity in order to propagate synchronous activity. However, for this to happen, the group size *M* must be much smaller than the network size *N^E^*. Here *N^E^* was fixed to 20,000 neurons for easier comparison of scenarios, but results are also valid for larger networks (see Materials and Methods). The good agreement between the mean-field theory and the numerical results suggests that the crucial parameter for assembly-sequence replay is the total input one neuron is receiving, e.g., the number of input synapses.

### Stronger synapses are equivalent to more connections

Up to this point, all excitatory synaptic connections in our model had constant and equal strengths. By encoding an assembly sequence we implicitly altered the structural connectivity by creating new synaptic connections. This case of structural plasticity can also occur when silent synapses are turned into functionally active connections upon learning [4, 38]. However, learning new associations might also be possible through a change of synaptic strength of individual connections [10, 64]. If a sequence is to be learned through synaptic plasticity, then instead of increasing the connectivity between groups of neurons, the synaptic conductances could be increased as well. To test whether these two types of plasticity are equivalent in our approach, we embed assembly sequences with various feedforward connectivities *p*_ff_ and various feedforward conductances 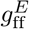, while keeping the recurrent connectivity (*p*_rc_ = 0.06) and recurrent conductances ( *g^E^* = 0.1 nS) constant.

Numerical results show that feedforward connectivity and feedforward conductance have identical roles in the replay of a sequence. That is, the sparser the connections, the stronger synapses are required for the propagation of activity. The analytical estimate (Fig 7A, black line corresponds to 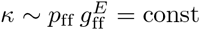.) predicts that the product of p _ff_and 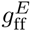 is the essential parameter for replay.

**Fig 7.**
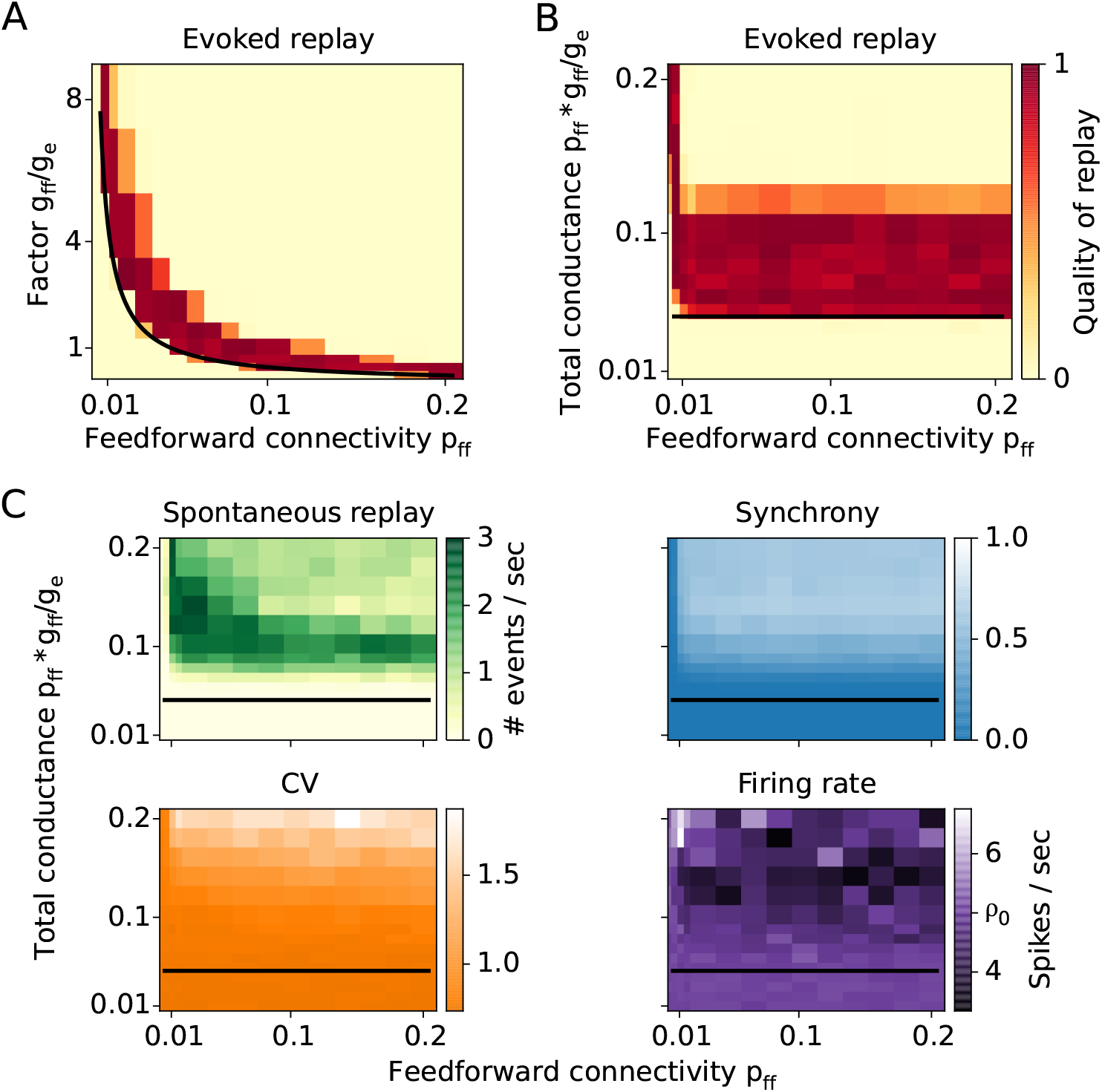
Feedforward conductance versus feedforward connectivity. **A:**Quality of replay as a function of connectivity and synaptic strength. **B:** The replay as a function of connectivity and total feedforward conductance input shows that the propagation is independent of connectivity as long as the total feed-forward input is kept constant. **C:** Spontaneous network dynamics described by the rate of spontaneous replay, synchrony, CV, and firing rate.

That this analytical prediction is fulfilled in the numerical simulations becomes clearer when we show the replay quality as a function of the feedforward connectivity and the total feedforward input 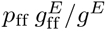 a neuron is receiving (Fig 7B). It is irrelevant whether the number of connections are changed or their strength, what matters is their product. This rule breaks only for sparse connectivities (*p*_ff_ < 0.01), i.e. when the mean number of feedforward connections between two groups is low ( <5). Therefore, the number of relevant connections cannot be reduced to very low numbers.

Consistent with earlier findings, the quality of replay is high above a certain strength of the total feedforward conductance (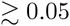 in Fig 5B) and for *p*_ff_ > 0.01. However, for sufficiently large feedforward input 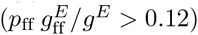, the replay of sequences is severely impaired as the network is in a state of highly synchronous bursting activity (Fig 7B), which is similar to the results shown in Figs 5 and 6.

The rule that the total input 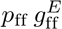 determines the network behavior also holds for spontaneous activity. Spontaneous replay rate, CV, synchrony, and firing rate all vary as a function of the total input (Fig 7C), and only weakly as a function of the connectivity or the conductance alone. Similar to the previous results in Figs 5 and 6, for 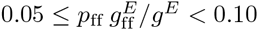 it is possible to evoke a replay while preserving the AI state of the network. Increasing the total input beyond this value drives the network into a state of spontaneous replay with increased synchrony.

### Forward and reverse replay in assembly sequences with symmetric connections

The assembly-sequence model discussed until now contains asymmetric connections, i.e., neurons from one group project extensively within the same and the subsequent group but not to the previous group. We showed that such feedforward assembly sequences are capable of propagating activity, which we call replay. Thus, the proposed model may give an insight on the replay of behavioral sequences that have been observed in the hippocampus [58]. However, further experiments revealed that sequences are also replayed in the inverse temporal order than during behavior, so-called reverse replay [23, 30]. The direction of this replay also depended on the context, i.e., when the animal was at the beginning of the path, forward replays prevailed; while after traversing the path, more reverse replays were detected (but see [47]). This suggests that the replay activity might be cued by the sensory input experienced at the current location of the animal.

As the feedforward structure adopted in the network model is largely asymmetric, the assembly sequence is incapable of reverse replay in its current form. To be able to activate a sequence in both directions, we modify the network and add symmetric connectivity between assemblies [62, 68, 82]. Then, an assembly of neurons does not project only to the subsequent assembly but also to the preceding, and both projections are random with probability *p*_ff_ (Fig 8A). While this connectivity pattern decreases the group clustering, it does not lead to full merge of the assemblies because the inhibition remains local for each group.

**Fig 8.**
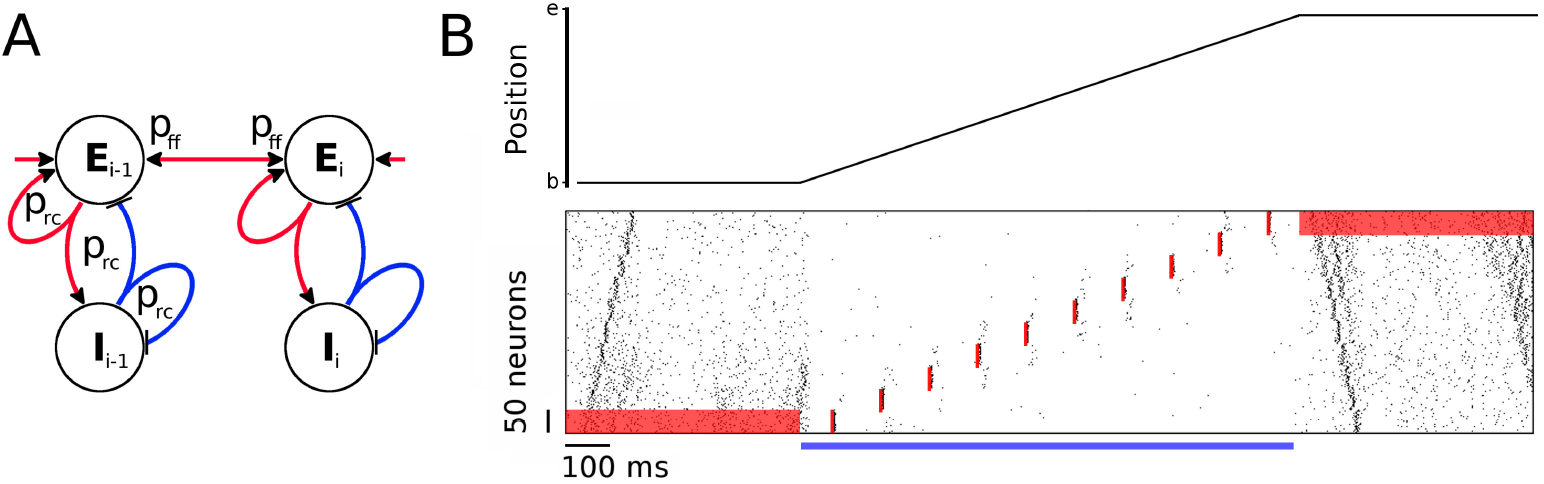
Symmetric assembly sequence. **A:**Schematic of an assembly sequence with symmetric connections between groups. **B:** Virtual rat position on a linear track (top) and the corresponding neuronal activity (bottom) as a function of time for 2 seconds. The rat rests at position “b” for half a second, then moves from “b” to “e” with constant speed for one second, where it rests for another 500 ms. While the rat is immobile at both ends of the track, a positive current input *I ^e^* = 2 pA is applied to the excitatory population of the first and last assembly as shown by the red background in the raster plot. Spontaneous replays start from the cued assemblies. During exploration, however, the network activity is decreased by a current *I^e^* = − 10 pA injected to the whole excitatory population, denoted with a blue horizontal bar. Strong sensory input during traversal activates the location-specific assemblies but does not result in any replay. The timing and location of the stimulations is denoted with red vertical bars in the raster plot. Recurrent and feedforward connectivities are *p*_rc_ = 0.15 and *p*_ff_ = 0.03, respectively.

Interpreting this network as a model for hippocampal activity during spatial navigation of a virtual rat on a linear track (Fig 8B, top), we test the idea that external input can switch the network between a spontaneous-replay state during rest and a non-replay, spatial-representation state during locomotion. During immobility at the beginning of the track, a context-dependent input cue is mimicked by a constant current *I ^e^* = 2 pA injected into the excitatory neurons of the first assembly (Fig 8B, red bar from 0 to 500 ms). The elevated firing rate of the first assembly results in a spontaneous forward replay, similar to the experimental findings during resting states at the beginning of a linear track [23, 30]. In contrast, in the absence of the context-dependent current, spontaneous replay can start at any assembly in the sequence (as in Fig 5) and propagate in forward or reverse direction.

After the initial 500 ms resting period, an external global current of − 10 pA is injected into the whole excitatory population to decrease network excitability and to mimic a state in which the rat explores the environment. In addition, to model place-specific sensory input that is locked to theta oscillations, we apply a strong and brief conductance input (as in Fig 2) every 100 ms to the assembly that represents the current location. In this situation, the assemblies fire at their corresponding locations only. There is, however, a weak activation of the neighboring assemblies that does not result in a replay. An extension of the model including lateral inhibition and STP would possibly enable theta sequences that span in one direction only [80]. Such an extension is, however, beyond the scope of the current manuscript.

At the end of the track, we retract the global external current to return to the virtual resting state for the last 500 ms of the simulation, and the network switches back to higher mean firing rates. A context-dependent sensory cue to the last group ( *I ^e^* = 2 pA current injected continuously) then leads to a spontaneous reverse replay, similar to experimental findings at the end of a linear track [23, 30].

In summary, we show that given symmetric connectivity between assemblies, transient activity can propagate in both directions. Large negative external currents injected into all excitatory neurons can decrease network excitability and thus block the replay of sequences. On the other hand, spontaneous replay can be cued by a small increase in the firing rate of a particular assembly. Interestingly, once a replay is initiated, it does not change direction, in spite of the symmetric connectivity. An active assembly receives feedback inhibition from its inhibitory subpopulation, which prevents immediate further activations and hence a reversal of the direction of propagation.

## Discussio

We revived Hebb’s idea on assembly sequences (or “phase sequences”) where activity in a recurrent neural network propagates through assemblies [41], a dynamics that could underlie the recall and consolidation of memories. An important question in this context is how learning of a series of events can achieve a strong enough synaptic footprint to replay this sequence later. Using both numerical simulations of recurrent spiking neural networks and an analytical approach, we provided a biologically plausible model for understanding how minute synaptic changes can nevertheless be uncovered by small cues or even manifest themselves as activity patterns that emerge spontaneously. We showed how the impact of small changes in the connections between assemblies is boosted by recurrent connectivity within assemblies. This interaction between recurrent amplification within an assembly and the feedforward propagation of activity establishes a possible basis for the retrieval of memories. Our theory thus provides an unifying framework that combines the fields of Hebbian assemblies and assembly sequences [41], synfire chains [1, 25], and fast amplification in balanced recurrent networks that are in an asynchronous-irregular state [72, 94].

Main conclusions from our work are that the effective coupling between assemblies is a function of both feedforward and recurrent connectivities, and that the network can express three main types of behavior: 1. When the coupling is weak enough, assembly sequences are virtually indistinguishable from the background random connections, and no replays take place. 2. For sufficiently strong coupling, a transient input to some assembly propagates through the sequence, resulting in a replay. 3. For even stronger coupling, noise fluctuations get amplified by the underlying structure, resulting in spontaneous replays. Each of these three regimes has a certain advantage in performing a particular task. Weak coupling is appropriate for imprinting new sequences if the network dynamics is driven by external inputs rather than controlled by the intrinsically generated activity. Intermediate coupling is suitable for recollection of saved memories; sequences remain concealed and are replayed only by specific input cues; otherwise, the network is in the asynchronous-irregular, spontaneous state. For strong coupling, spontaneous replays might be useful for offline recollection of stored sequences when there are no external input cues. Importantly, the network behaviour and the rate of spontaneous events depends not only on the coupling but can be controlled by modulating the network excitability through external input. Neuromodulator systems, for example the cholinergic and the adrenergic systems [40, 87] might therefore mediate the retrieval process.

In our simulations, we examined relatively short and non-overlapping assemblies. A natural question is whether the network can sustain longer chains and tolerate overlapping patterns. In additional simulations (results not shown), we found that both is possible. However, because previous work has dealt in great detail with the calculation of capacity of neural networks both analytically [59, 60] and computationally [89], we did not explore this issue.

### Related models

Assembly sequences are tightly related to synfire chains, which were proposed [1] as a model for the propagation of synchronous activity between groups of neurons. Diesmann et al. [25] showed for the first time that synfire chains in a noisy network of spiking neurons can indeed support a temporal code. It has been shown, however, that the embedding of synfire chains in recurrent networks is fragile [6, 66], because on the one hand, synfire chains require a minimal connectivity to allow propagation, while on the other hand, a dense connectivity between groups of neurons can generate unstable network dynamics. Therefore, Aviel et al. [5] introduced “shadow pools”of inhibitory neurons that stabilize the network dynamics for high connectivity. The network fragility can also be mitigated by reducing the required feedforward connectivity: inputs from the previous assembly are boosted by recurrent connections within the assembly. This approach was followed Kumar et al. [54], which examined synfire chains embedded in random networks with local connectivity, thus, implicitly adopting some recurrent connectivity within assemblies as proposed by the assembly-sequence hypothesis; nevertheless, their assemblies were fully connected in a feedforward manner. Recently, it was shown that replay of synfire chains can be facilitated by adding feedback connections to preceding groups [69]. However, this Hebbian amplification significantly increased the duration of the spike volleys and thus decreased the speed of replay. Our model circumvents this slowing effect by combining the recurrent excitation with local feedback inhibition, effectively replacing Hebbian amplification by a transient “balanced amplification” [72].

To store sequences, further classes of models were proposed, e.g., “winner-takes-all” [46, 50, 71] and “communication through resonance” [37]. However, the activity propagation in these models has an order of magnitude slower time scales than the synfire chain or the assembly sequence, and thus, are not suitable for rapid transient replays.

The spontaneous replay in our network bears some resemblance with the population bursts that occur in a model with supralinear amplification of precisely synchronised inputs [67]. Adding such nonlinearities to the conductances in our model might decrease even further the connectivity required for the assembly-sequence replay. Another model class, which relies on lognormal conductance distributions, has been proposed as a burst generator for SWRs [74]. The model accounts for spontaneously generated stereotypical activity that propagates through neurons that are connected with strong synapses.

To summarize, for the propagation of activity, functionally connecting assemblies of excitatory and inhibitory neurons requires lower number of additional feedforward synapses than connecting random groups of neurons. This lower number of synapses may facilitate rapid, single-shot learning of associations and enhance the memory capacity of the network [59, 90].

### Relation between recurrent and feedforward connectivity

What is the most efficient set of connectivities in terms of numbers of synapses used? To create an assembly of *M* neurons and to connect it to another assembly of the same size, we need *M*^2^(*p*_rc_ + *p*_ff_) excitatory-to-excitatory synapses. The constraint *к* = 1 then leads to a minimum total number of synapses at *p*_rc_ = 0. This result is somewhat surprising because it suggests that our proposed recurrent amplification provides a disadvantage.

However, another constraint might be even more important: to imprint an association in one-shot learning, as for example required for episodic memories, it might be an advantage to change as few synapses as possible so that one can retrieve the memory later via a replay. Therefore, *p*_ff_ should be low, in particular lower than the recurrent connectivity that is bound by the morphological connectivity that includes also weak or silent synapses. Minimizing *p*_ff_ under the constraint *к* = 1 implies, however, maximizing *p*_rc_. Such large connectivities might require longer time to develop. A large *p*_rc_ is compatible with one-shot learning only if assemblies (that are defined by increased *p*_rc_ among a group of neurons) can be set up *prior* to the (feedforward) association of assemblies. Thus, episodic memories could benefit from strong preexisting assemblies. For setting up such assemblies, long time periods might be available to create new synapses and to morphologically grow synapses. Thus, we predict that for any episodic memory to be stored in one shot learning in hippocampal networks such as CA3, a sufficiently strong representation of the events to be associated does exist *prior* to successful one-shot learning. In this case, *p*_ff_ (i.e., connectivity in addition to *p*_rand_) can be almost arbitrarily low. A natural lower limit is that the number of synapses per neuron *M _pff_* is much larger than 1, say 10 as a rough estimate (in Fig 3 we have *M*_pff_ ∼ 30 for a rather low value of *p*_rc_ = *p*_ff_, and 10 for *p*_rc_ = 0.30; even 5 or more very strong synapses are sufficient in Fig 7), which can be interpreted in two ways: (1) Every neuron in an assembly should activate several neurons in the subsequent assembly, and (2) every neuron in an assembly to be activated should receive several synapses from from neurons in the previous assembly.

For example in the modeled network, for *p*_ff_ = 0.02 and *M _pff_* > 10 we obtain *M >* 500, which is in agreement with an estimated optimal size of assemblies in the hippocampus [59]. The total number of feedforward synapses required for imprinting an association is then *M*^2^ *p*_ff_ > 5,000, which is a relatively small number compared to the total number of background synapses 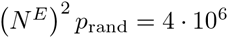 for *N^E^* = 20,000 and *p*_rand_ = 0.01. Scaling up the network accordingly (see Materials and Methods) to the size of a mouse CA3 network, i.e., *N ^E^* = 240,000 (a typical number for the rat hippocampus, e.g., [76, 98]), the number of new associative synapses is *M*^2^*p*_ff_ > 17,000, while the total connections are more than 0.5.10^9^.

To conclude, abundant recurrent connections within assemblies can decrease the feedforward connectivity required for a replay to almost arbitrary low values. Moreover, the ratio of memory synapses to background synapses decreases as the network is scaled to bigger size.

### Mechanisms for assembly-sequence formation

For sequence replay, increasing the number of connections between groups has the same effect as scaling up the individual connection strengths. We conclude that structural and synaptic plasticity could play an equivalent role in the formation of assembly sequences. However, in the current study we have not considered plasticity mechanisms that could be mediating the formation of assembly sequences. Previous attempts of implementing a spike-timing-dependent plasticity (STDP) rule with an asymmetric temporal window [9, 32, 48] in recurrent networks led to structural instabilities [43, 57, 70]. More sophisticated learning rules better matched the experimentally observed plasticity protocols [17, 35, 75], and these rules combined with various homeostatic mechanisms could form Hebbian assemblies that remained stable over long time periods [62, 82, 102]. Moreover, [62] and [82] have shown that the voltage-based STDP rule [17] leads to strong bidirectional connections, a network motif that has been reported in multiple brain regions [20, 51, 83, 85]. A recent experimental work on the plasticity of the CA3-to-CA3 pyramidal cell synapses has revealed a symmetric STDP temporal curve [68]. Such a plasticity rule can be responsible for the encoding of stable assembly representations in the hippocampus.

Several plasticity rules have been successfully applied in learning sequences [11, 53, 78, 84, 95]. However, these studies focused purely on sequence replay and did not take into account its interaction with a balanced, asynchronous irregular background state.

### Relations to hippocampal replay of behavioral sequences

The present model may explain the replay of sequences associated with the sharp-wave ripple (SWR) events, which originate in the CA3 region of the hippocampus predominantly during rest and sleep [14]. SWRs are characterized by a massive neuronal depolarization reflected in the local field potential [21]. Moreover, during SWRs, pyramidal cells in the CA areas fire in sequences that reflect their firing during prior awake experience [58]. Cells can fire in the same or in the reverse sequential order, which we refer to as forward and reverse replay, respectively [23, 30].

According to the two-stage model of memory trace formation [14], the hippocampus is encoding new episodic memories during active wakefulness (stage one). Later, these memories are gradually consolidated into neocortex through SWR-associated replays (stage two). It has been proposed that acetylcholine (ACh) modulates the flow of information between the hippocampus and the neocortex and thereby mediates switches between these memory-formation stages [39]. During active wakefulness, the concentration of ACh in hippocampus is high, leading to partial suppression of excitatory glutamatergic transmission [40] and promoting synaptic plasticity [36]. In this state, a single experience seems to be sufficient to encode representations of the immediate future in an environment [29]. On the other hand, the level of ACh decreases significantly during slow-wave sleep [65], releasing the synaptic suppression and resulting in strong excitatory feedback synapses, which suggests that this boost of recurrent and feedback connections leads to the occurrence of SWRs. In line with this hypothesis, the present model shows that increasing the synaptic strengths shifts the assembly-sequence dynamics from a no-replay regime to a spontaneous-replay regime. Also, we demonstrated that this regime supports both forward and reverse replay if assemblies are projecting symmetrically to each other and if recurrent connectivity exceeds severalfold the feedforward coupling.

In summary, a prediction of our assembly-sequence model is that prior to being able to store and recall a memory trace that connects events, strong enough representations of events in recurrently connected assemblies are necessary because recalling a minute memory trace requires amplification within assemblies. Another prediction of this model is based on the fact that the network is in an asynchronous-irregular state during the time intervals between replays. Hence, by increasing the activity of the excitatory neurons or by disinhibiting the network, e.g., by decreasing the activity of the interneuron population specialized in keeping the balance, one could increase the rate of spontaneous replays. Our model thus links a diverse set of experimental results on the cellular, behavioral, and systems level of neuroscience on memory retrieval and consolidation [24].

## Materials and Methods

The network simulations as well as the data analyses were performed in Python (www.python.org). The neural network was implemented in Brian [34]. For managing the simulation environment and data processing, we used standard Python libraries such as NumPy, SciPy, Matplotlib, and SymPy.

### Neuron model

Neurons are described by a conductance-based leaky integrate-and-fire model, where the subthreshold membrane potential *Vi*(*t*) of cell *i* obeys

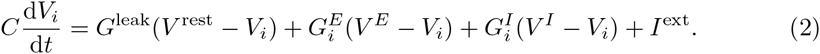

The cells’ resting potential is *V*^rest^ = −60 mV, its capacitance is *C* = 200 pF, and the leak conductance is *G*^leak^ = 10 nS, resulting in a membrane time constant of 20 ms in the absence of synaptic stimulation. The variable 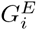 and 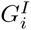 are the total synaptic conductances describing the time-dependent synaptic inputs to neuron *i*. The excitatory and inhibitory reversal potentials are *V ^E^* = 0 mV and *V ^I^* = − 80 mV, respectively. *I*^ext^ = *I*^const^ + *I^x^* is an externally applied current. To evoke activity in the network, a constant external current *I*^const^ = 200 pA is injected into each neuron. Only if explicitly stated (e.g., Figs 5 and 8), small additional current inputs *I ^x^* are applied to excitatory or inhibitory neurons, which we denote as *I^e^* and *I^i^*, respectively. As the membrane potential *V_i_* reaches the threshold *V*^th^ = −50 mV, neuron *i* emits an action potential, and the membrane potential *Vi* is reset to the resting potential *V*^rest^ for a refractory period *τ*_rp_ = 2 ms.

The dynamics of the conductances 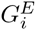 and 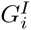 of a postsynaptic cell *i* are determined by the spiking of the excitatory and inhibitory presynaptic neurons. Each time a presynaptic cell *j* fires, the synaptic input conductance of the postsynaptic cell *I* is increased by 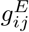 for excitatory synapses and by 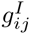 for inhibitory synapses. The input conductances decay exponentially with time constants *τ^E^* = 5 ms and *τ^I^* = 10 ms. The dynamics of the total excitatory conductance is described by

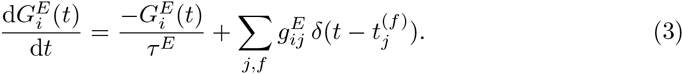

Here the sum runs over the presynaptic projections *j* and over the sequence of spikes *f* from each projection. The time of the *f*^th^ spike from neuron *j* is denoted by 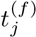, and *δ* is the Dirac delta function. The inhibitory conductance 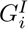 is described analogously.

Amplitudes of recurrent excitatory conductances and excitatory conductances on inhibitory neurons are denoted with 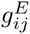 and 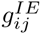, respectively. If not stated otherwise, all excitatory conductance amplitudes are fixed and equal 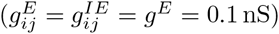, which results in EPSPs with an amplitude of ≈ 0.1 mV at resting potential. The recurrent inhibitory synapses are also constant 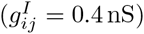 while the inhibitory-to-excitatory conductances 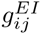 are variable (see below). Irrespectively of the synaptic type, the delay between a presynaptic spike and a postsynaptic response onset is always 2 ms.

### Network model

The modelled network consists of *N ^E^* = 20,000 excitatory and *N ^I^* = 5,000 inhibitory neurons. Our results do not critically depend on the network size (see Section ‘Scaling the network size’ below). Initially, all neurons are randomly connected with a sparse probability *p*_rand_ = 0.01.

A cell assembly is defined as a group of recurrently connected excitatory and inhibitory neurons (Fig 1A). The assembly is formed by picking *M* excitatory and *M/*4 inhibitory neurons from the network; every pair of pre- and post-synaptic neurons within the assembly is randomly connected with probability *p*_rc_. The new connections are created independently and in addition to the already existing ones. Thus, if by chance two neurons have a connection due to the background connectivity and are connected due to the participation in an assembly, then the synaptic weight between them is simply doubled. Unless stated otherwise, assemblies are hence formed by additional connections rather than stronger synapses.

In the random network, we embed 10 non-overlapping assemblies with size *M* = 500 if not stated otherwise. The groups of excitatory neurons are connected in a feedforward fashion, and a neuron from one group projects to a neuron of the subsequent group with probability *p*_ff_ (Fig 1B). Such a feedforward connectivity is reminiscent of a synfire chain. However, classical synfire chains do not have recurrent connections (*p*_rc_ = 0, *p*_ff_ > 0), while here, neurons within a group are recurrently connected even beyond the random background connectivity (*p*_rc_ > 0, *p*_ff_ > 0). We will refer to such a sequence as an “assembly sequence”. By varying the connectivity parameters *p* _rc_and *p*_ff_, the network structure can be manipulated to obtain different network types (Fig 1C). In the limiting case where feedforward connections are absent (*p*_rc_ > 0, *p*_ff_ = 0) the network contains only largely disconnected Hebbian assemblies. In contrast, in the absence of recurrent connections (*p*_rc_ = 0, *p*_ff_ > 0), the model is reduced to a synfire chain embedded in a recurrent network. Structures with both recurrent and feedforward connections correspond to Hebbian assembly sequences.

To keep the network structure as simple as possible and to be able to focus on mechanisms underlying replay, we use non-overlapping assemblies and we do not embed more than 10 groups. Nevertheless, additional simulations with overlapping assemblies and longer sequences (results not shown) indicate that our approach is in line with previous results on memory capacity [59, 60, 89]. Advancing the theory of memory capacity is, however, beyond the scope of this manuscript.

## Balancing the network

A naive implementation of the heterogeneous network as described above leads, in general, to dynamics characterized by large population bursts of activity. To overcome this epileptiform activity and ensure that neurons fire asynchronously and irregularly (AI network state), the network should operate in a balanced regime. In the balanced state, large excitatory currents are compensated by large inhibitory currents, as shown *in vivo* [15, 73] and *in vitro*[101]. In this regime, fluctuations of the input lead to highly irregular firing [92, 93].

Several mechanisms were proposed to balance numerically simulated neural networks. One method involves structurally modifying the network connectivity to ensure that neurons receive balanced excitatory and inhibitory inputs [77, 81]. It was shown that a short-term plasticity rule [91] in a fully connected network can also adjust the irregularity of neuronal firing [8].

Here, we balance the network using the inhibitory-plasticity rule [94]. All inhibitory-to-excitatory synapses are subject to a spike-timing-dependent plasticity (STDP) rule where near-coincident pre- and postsynaptic firing potentates the inhibitory synapse while presynaptic spikes alone cause depression. A similar STDP rule with a symmetric temporal window was recently reported in the layer 5 of the auditory cortex [22].

To implement the plasticity rule in a synapse, we first assign a synaptic trace variable *x_i_* to every neuron *i* such that *x_i_* is incremented with each spike of the neuron and decays with a time constant *τ*_STDP_ = 20 ms:

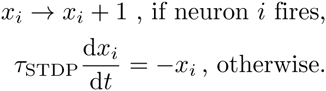

The synaptic conductance 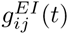 from inhibitory neuron *j* to excitatory neuron *i* is initialized with value 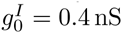 and is updated at the times of pre/post-synaptic events:

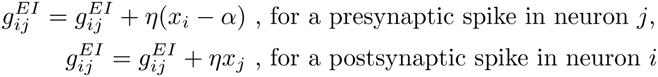

where 0 < *η* ≪ 1 is the learning-rate parameter, and the bias *α* = 2*ρ*_0_ *τ*_STDP_ is determined by the desired firing rate *ρ*_0_ of the excitatory postsynaptic neurons. In all simulations, *ρ*_0_ has been set to 5 spikes/sec, which is at the upper bound of the wide range of rates that were reported in the literature: e.g., 1 −3 spikes/sec in [21]; 3 −6 spikes/sec in [52]; 1 − 76 spikes/sec in [28]; 0.43 −3.60 spikes/sec in [16]; 1 − 11 spikes/sec in [27].

An implementation of the described STDP rule drives typically the network into a balanced state. The excitatory and the inhibitory input currents balance each other and keep the membrane potential just below threshold while random fluctuations drive the firing (Fig 2A,B). The specific conditions to be met for a successful balance are discussed in the Results section.

In the AI network regime, any perturbation to the input of an assembly will lead to a transient perturbation in the firing rate of the neurons within it. Moreover, because of the recurrent connections, even small perturbations can lead to large responses. This phenomenon of transient pattern completion is known as balanced amplification [72], where it is essential that each assembly has excitatory and inhibitory neurons. Another advantage of the inhibitory subpopulations is the rapid negative feedback that can lead to enhanced memory capacity of the network [45].

### Simulations and data analysis

Each network simulation consists of 3 main phases:

#### 1. Balancing the network

Initially, the population activity is characterized by massive population bursts with varying sizes (avalanches). During a first phase, the network (random network with embedded phase sequence) is balanced for 50 seconds with decreasing learning rate (0.005 ≥ η ≥ 0.00001) for the plasticity on the inhibitory-to-excitatory synapses. During this learning, the inhibitory plasticity shapes the activity, finally leading to AI firing of the excitatory population. Individual excitatory neurons then fire roughly with the target firing rate of 5 spikes/sec, while inhibitory neurons have higher firing rates of around 20 spikes/sec, which is close to rates reported in the hippocampus [16, 21]. After 50 seconds simulation time, the network is typically balanced.

#### 2. Reliability and quality of replay

In a second phase, the plasticity is switched off to be able to probe an unchanging network with external cue stimulations. All neurons from the first group/assembly are simultaneously stimulated by an external input so that all neurons fire once. The stimulation is mimicked by adding an excitatory conductance in Eq 3 (^g_max_^ = 3 nS) that is sufficient to evoke a spike in each neuron. For large enough connectivities (*p*_rc_ and *p*_ff_), the generated pulse packet of activity propagates through the sequence of assemblies, resulting in a replay. For too small connectivities, the activity does not propagate. For excessively high connectivities, the transient response of one group results in a burst in the next group and even larger responses in the subsequent groups, finally leading to epileptiform population bursts of activity (Fig 3).

To quantify the propagation from group to group and to account for abnormal activity, we introduce a quality measure of replay. The activity of a group is measured by calculating the population firing rate of the underlying neurons smoothed with a Gaussian window of 2 ms width. We extract peaks of the smoothed firing rate that exceed a threshold of 30 spikes/sec. A group is considered to be activated at the time at which its population firing rate hits its maximum and is above the threshold rate. Activity propagation from one group to the next is considered to be successful if one group activates the next one within a delay between 2 and 20 ms. A typical delay is about 5 ms, but in the case of extremely small *p*_ff_ and large *p*_rc_ the time of propagation can take ∼ 15 ms. Additional rules are imposed to account for exceeding activity and punish replays that lead to run-away firing. First, if the activity of an assembly exceeds a threshold of 180 spikes/sec (value is chosen manually for best discrimination), the group is considered as bursting, and thus, the replay is considered as failed. Second, if the assembly activity displays 2 super-threshold peaks that succeed each other within 30 ms, the replay is unsuccessful. Third, a “dummy group” (of size *M*) from the background neurons is used as a proxy for detecting activations of the whole network. In case that the dummy group is activated during an otherwise successful replay, the replay is failed. Thus, for each stimulation the “quality of replay” has a value of 1 for successful and a value of 0 for unsuccessful replays. The quality of replay for each set of parameters (Fig 3) is an average from multiple 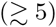 stimulations of 5 different realizations of each network.

Additionally, we test the ability of the assembly sequence to complete a pattern by stimulating only a fraction of the neurons in the first group (Fig 4). Analogously to the full stimulation, the quality of replay is measured.

#### 3. Spontaneous activity

In the last phase of the simulations, no specific input is applied to the assemblies. As during the first phase of the simulation, the network is driven solely by the constant-current input *I*^const^ = 200 pA applied to each neuron, and plasticity is switched off.

During this state, we quantify spontaneous replay (Fig 5). Whenever the last assembly is activated and if this activation has propagated through at least three previous assemblies, we consider this event as a spontaneous replay. Here, we apply the quality measure of replay, where bursty replays are disregarded. Additionally, we quantify the dynamic state of the network by the firing rate, the irregularity of firing, and the synchrony of a few selected groups from the sequence. The irregularity is measured as the average coefficient of variation of inter-spike intervals of the neurons within a group. As a measure of synchrony between 2 neurons, we use the cross-correlation coefficient of their spike trains binned in 5-ms windows. The group synchrony is the average synchrony between all pairs of neurons in a group.

### Mean-field analysis

To analytically describe the conditions for a successful sequence replay, we portray the network activity during replay using a linear model. Approximating the network dynamics with a system of linear differential equations, we estimate a lower bound for the connectivities required for a successful replay.

The dynamics of an assembly *i* (Fig 1A, B) in the AI state is approximated by two differential equations:

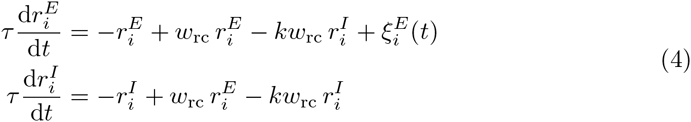

where 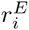 and 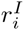are the deviations of the population firing rates of the excitatory (E) and inhibitory (I) populations from the spontaneous firing rates 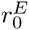 and 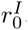, respectively.

The parameter *w*_rc_ and the term *−kw*_rc_ are the respective strengths of the excitatory and the inhibitory recurrent projections. The constant *k* describes the relative strength of the recurrent inhibition vs. excitation; for a balanced network, we assume that inhibition balances or dominates excitation, e.g., *k* ≥ 1. The weight *w*_rc_ is proportional to the average number *M p*_rc_ of recurrent synapses a neuron receives, and proportional to the synaptic strength *g^E^*. The function 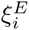 describes the external input to the assembly from the rest of the network. In this mean-field analysis, we neglect the influence of the noise on the network dynamics. Activities 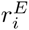 and 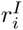 are assumed to approach the steady state 0 with a time constant *τ*.

The excitatory assemblies are sequentially connected, and we denote the strength of the feedforward projections as *w*_ff_. The feedforward drive can be represented as an external input to an assembly:

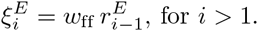

Taking into account the feedforward input to population *i* from the preceding excitatory population *i −*1, Eq 4 can be rewritten as

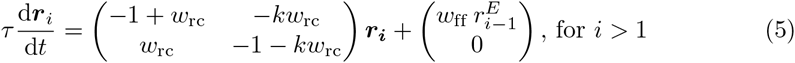

where 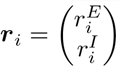 the 2-dimensional vector of firing rates in group *i*.

From previous theoretical studies [31, 92, 93] we know that in the AI state the population time constant *τ* can be much smaller than the membrane time constant of individual neurons. This means that a population of neurons can react faster to external input than individual neurons. Assuming that the time duration of a pulse packet in group *i −*1 is much longer than the population time constant *τ*in group *i*, wecan consider the solution of the stationary state 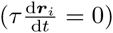 as an adequate approximation. Thus, by setting the left-hand side of Eq 5 to zero, we can express the firing rate 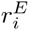 as a function of 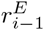

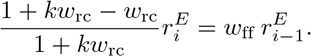

In the special case of a balanced network where *k* = 1, the relation can be simplified further:

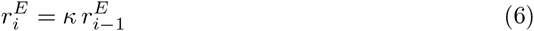

where

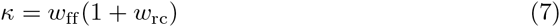

is the “effective feedforward connectivity”. Interestingly, the recurrent connections effectively scale up the efficiency of the feedforward connections and facilitate the propagation of activity. For small *к*, i.e. *к* ≪ 1, even large changes of the firing rate in group *i –*1 do not alter the rate in group *i*. For *к* < 1, the pulse packet will steadily decrease while propagating from one group to another as 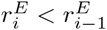 On the other hand, if к = 1, the propagation of a pulse packet is expected to be stable. In the case of *к* >1, any fluctuation of firing rate in one assembly will lead to a larger fluctuation in the following assembly.

To connect the analytical calculations to the numerical simulations, we again note that a total connection strength is proportional to the number of inputs a neuron is receiving (e.g., the product of group size *M* and connection probability) and proportional to the synaptic strength:

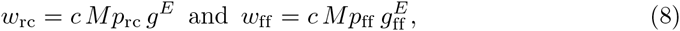

where *M* is the group size, and *p*_rc_ and *p*_ff_ are the recurrent and feedforward connectivities, respectively. *g^E^*is the conductance of an excitatory recurrent synapse within a group, and 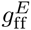 is the conductance of feedforward synapses between groups. Unless stated otherwise, we assume 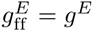 The parameter *c* is related to the slope of the neurons’ input-output transfer function.

Before representing *к* as function of the connectivities *p*_ff_ and *p*_rc_, we estimate the parameter *c*. By fitting the critical value *к*(*p*_rc_ = 0.08, *p*_ff_ = 0.04) = 1 from the simulation results (Fig 3), we find *c* = 0.25 nS^−1^. This value of *c* is used in all further analytical estimations for the effective connectivity *к*. However, this procedure does not show us how *c* depends on various parameters, e.g., conductances, time constants, network size, etc. Therefore, the next subsection deals with deriving an explicit expression for the transfer function slope *c*.

In summary, the lower bound for the connectivities for a successful replay can be described as

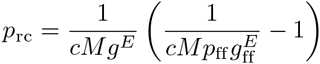

which is represented as a black line in Figs 3 and 5. For Figs 6 and 7, the black line is calculated analogously using the same constant *c* = 0.25 nS^−1^.

### Calculating the slope *c*

In the previous section, the constant *c* was manually fitted to a value of 0.25 nS^−1^ to match analytical and numerical results. Here we express *c* analytically by utilizing a non-linear neuronal model and by using the parameter values from the simulations.

The resting firing rate *ρ* of a neuronal population that is in an asynchronous irregular (AI) regime can be expressed as a function of the mean *µ* and the standard deviation *σ* of the membrane potential distribution [3, 13, 33, 79]:

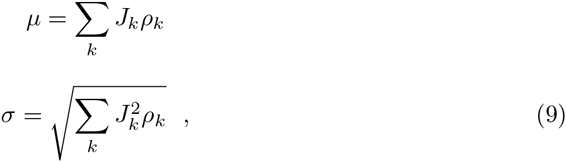

where the sums over *k* run over the different synaptic contributions, *ρ_k_* is the corresponding presynaptic firing rate, and *J_k_* and 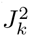 are the integrals over time of the PSP and the square of the PSP from input *k*, respectively. Here PSPs are estimated for the conductance-based integrate-and-fire neuron from Eq 2 for voltage values near the firing threshold *V*^th^,

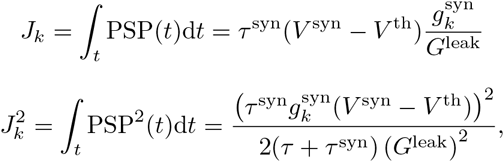

where *τ* is the membrane time constant, *τ^syn^* is the synaptic time constant, *V^syn^* is the synaptic reversal potential, and 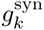 is the synaptic conductance of connection *k*. Connections can be either excitatory or inhibitory.

Here we consider a network with random connections only, and look at a subpopulation of size *M*, where *M ≪ N^E^*. For a more convenient analytical treatment, the recurrent connections within the group are neglected. This assumption does not affect the estimation of the transfer function slope, as *c* is independent on the type of inputs. The firing rate-fluctuations of the neuronal group are calculated as in Eq 6:

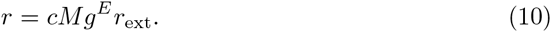

The membrane potential of an excitatory neuron from this subpopulation has several contributions: *N ^E^p_rand_* excitatory inputs with firing rate *ρ_0_* and efficacy *J^E^*; inhibitory inputs due to the background connectivity: 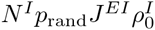 injected constant current: 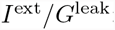; and input from an external group: 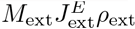. In summary, we find:

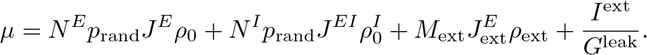

The standard deviation of the membrane potential is then, accordingly:

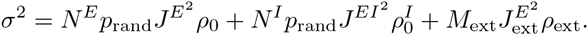

In the case of uncorrelated inputs, the following approximation can be used for the firing rate estimation [3, 13, 33, 79]:

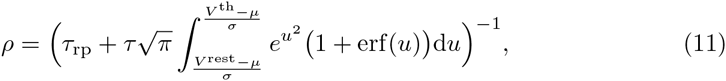

where *τ_rp_* is the refractory period, and *V*^th^and *V*^rest^ are membrane threshold and reset potential, respectively (see also section “Neural Model”).

To find the constant *c* used in the linear model, we estimate the firing rate *ρ* from Eq 11 and substitute in Eq 10, assuming a linear relation between firing-rate fluctuations:

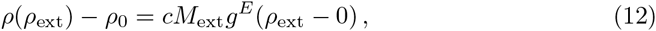

and find:

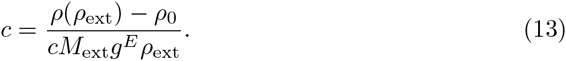

Before calculating the constant *c* according to the method presented above, a preliminary step needs to be taken. As we set the firing rate of the excitatory population in the network to a fixed value *ρ*_0_ = 5 spikes/sec, there are two variables remaining unknown: the firing rate of the inhibitory population 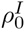 and the inhibitory-to-excitatory synaptic conductance 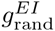 that changes due to synaptic plasticity. Therefore, we first solve a system of 2 equations for the firing rates of the excitatory and the inhibitory populations expressed as in Eq 11. Once the unknowns 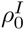 and 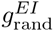 are calculated, we can estimate *ρ*(*ρ_ext_*) and *c* according to the method presented above. We note that the analytically calculated values of 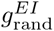 and 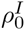 match the measured values in the simulations.

The value we get after applying the above mentioned method for estimation of *c* is 0.13 nS^−1^. The fit corresponding to the estimate of *c* is shown in Fig 3 with a white dashed line. It is worth noting that a slightly more involved calculation relying on the estimate 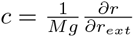 gives a similar result, concretely *c* = 0.11 nS^−1^.

Although the analytically calculated value *c* is a factor of 2 smaller than the manual fit *c* = 0.25 nS^−1^, it is qualitatively similar and not too far from describing the results for critical connectivity from the simulations.

The method applied above finds the slope of the transfer function for stationary firing rates. However, the spiking network replay is a fast and brief event, where a transient input in one assembly evokes a transient change in the output firing rate. The value discrepancy suggests that the transfer function of transients is even steeper than at the resting AI state.

### Scaling the network size

So far we have been dealing with networks of fixed size *N^E^* = 20,000 neurons. How does the network size affect the embedding of assembly sequences? Is it possible to change the network size but keep the assembly size fixed?

Scaling the network size while keeping the connectivity *p_rand_* constant leads to a change in the number of inputs that a neuron receives, and therefore, affects the membrane potential distributions. To compare replays in networks with different sizes *N ^E^* but identical *M*, we need to assure that the signal-to-noise ratio is kept constant, and the easiest way is to keep both the signal and the noise constant, which requires to change connectivities *p*_rc_ and *p*_ff_ and conductances.

While scaling the network from the default network size *N^E^* = 20,000 to a size 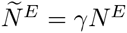, we see that the noise *σ* scales as 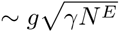 (Eq 9). To keep the input current fluctuations constant as we change 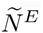, all synaptic conductances are rescaled with a factor of 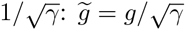 [92]. However, such a synaptic scaling leads to a change in the coupling between assemblies of fixed size *M*, which is proportional to the conductance. Therefore, the connectivities *p*_rc_ and *p*_ff_ are scaled with 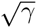 to compensate the conductance decrease, leading to a constant coupling 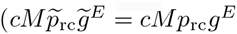, and 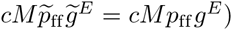 hence, a constant signal-to-noise ratio.

What is the impact of such a scaling on the network capacity to store sequences? The number of connections needed to store a sequence is changed by a factor 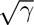 as we change *p*_rc_ and *p* _ff_. However, the number of background connections to each neuron is scaled with 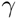, resulting in sparser memory representations in larger networks. More precisely, for a neuron participating in the sequence, the ratio of excitatory memory connections to the total number of excitatory connections is

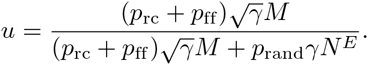

Therefore, the proportion of connections needed for an association is scaled as 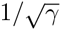 for 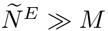 To give a few numbers, *u* is equal to 0.23 for 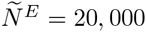, and *u* = 0.09 for.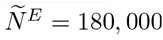 Other parameter values are: *M* = 500, *p*_rc_ = *p*_ff_ = 0.06, *p_rand_* = 0.01.

The chosen scaling rule is applicable for networks of simpler units such as binary neurons or current-based integrate-and-fire neurons [3, 93]. This scaling is not valid in a strict mathematical framework for very large networks 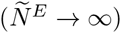 consisting of conductance-based integrate-and-fire neurons (see [77] for a detailed discussion).Simulations results, however, reveal that replays are possible in network sizes up to 2 · 10^5^neurons (results not shown).

## Acknowledgements

The authors thank José Donoso, André Holzbecher, and Jorge Jaramillo for insightful discussions and comments on the manuscript.

## References

1. Abeles M. Corticonics: neural circuits of the cerebral cortex. Cambridge, UK: Cambridge UP.; 1991.

2. Almeida-Filho DG, Lopes-dos-Santos V, Vasconcelos NA, Miranda JG, Tort AB, Ribeiro S. An investigation of Hebbian phase sequences as assembly graphs. Front Neural Circuits. 2014;8:34.

3. Amit DJ, Brunel N. Model of global spontaneous activity and local structured activity during delay periods in the cerebral cortex. Cereb Cortex.1997;7:,237–252.

4. Atwood HL, Wojtowicz JM. Silent synapses in neural plasticity: current evidence. Learn Mem. 1999;6:542–571.

5. Aviel Y, Horn D, Abeles M. Synfire waves in small balanced networks. Neurocomput. 2004;58:123–127.

6. Aviel Y, Mehring C, Abeles M, Horn D. On embedding synfire chains in a balanced network. Neural Comput. 2003;15:1321–1340.

7. Aviel Y, Pavlov E, Abeles M, Horn D. Synfire chain in a balanced network. Neurocomput. 2002;44:285–292.

8. Barbieri F, Brunel N. Can attractor network models account for the statistics of firing during persistent activity in prefrontal cortex? Front Neurosci. 2008;2:114–122.

9. Bi GQ, Poo MM. Synaptic modifications in cultured hippocampal neurons: dependence on spike timing, synaptic strength, and postsynaptic cell type. J Neurosci. 1998;18:10464–10472.

10. Bliss TV, Lømo T. Long-lasting potentiation of synaptic transmission in the dentate area of the anaesthetized rabbit following stimulation of the perforant path. J Physiol. 1973;232:331–356.

11. Brea J, Senn W, Pfister JP. Matching recall and storage in sequence learning with spiking neural networks. J Neurosci. 2013;33:9565–9575.

12. Brown GT. On the nature of the fundamental activity of the nervous centre: together with an analysis of the conditioning of rhythmic activity in progression, and a theory of the evolution of function in the nervous systems. J Physiol. 1914;48:18–46.

13. Brunel N. Dynamics of sparsely connected networks of excitatory and inhibitory spiking neurons. J Comput Neurosci. 2000;8:183–208.

14. Buzsaéki G. Two-stage model of memory trace formation: a role for “noisy” brain states. Neurosci. 1989;31:551–570.

15. Cafaro J, Rieke F. Noise correlations improve response fidelity and stimulus encoding. Nature. 2010;468:964–967.

16. Cheng J, Ji D. Rigid firing sequences undermine spatial memory codes in a neurodegenerative mouse model. eLife. 2013;2:e00647.

17. Clopath C, Buäsing L, Vasilaki E, Gerstner W. Connectivity reflects coding: a model of voltage-based STDP with homeostasis. Nat Neurosci. 2010;13:344–352.

18. Contreras EJB, Schjetnan AGP, Muhammad A, Bartho P, McNaughton BL, Kolb B, Gruber AJ, Luczak A. Formation and reverberation of sequential neural activity patterns evoked by sensory stimulation are enhanced during cortical desynchronization. Neuron. 2013;79:555–566.

19. Cossart R, Aronov D, Yuste R. Attractor dynamics of network UP states in the neocortex. Nature. 2003;423:283–288.

20. Cossell L, Iacaruso MF, Muir DR, Houlton R, Sader EN, Ko H, Hofer SB, Mrsic-Flogel TD. Functional organization of excitatory synaptic strength in primary visual cortex. Nature. 2015;518:399–403.

21. Csicsvari J, Hirase H, Mamiya A, Buzsaéki G. Ensemble patterns of hippocampal CA3-CA1 neurons during sharp wave-associated population events. Neuron. 2000;28:585–594.

22. D’amour JA, Froemke RC. Inhibitory and excitatory spike-timing-dependent plasticity in the auditory cortex. Neuron. 2015;86:514–528.

23. Diba K, Buzsaéki G. Forward and reverse hippocampal place-cell sequences during ripples. Nat Neurosci. 2007;10:1241–1242.

24. Diekelmann S, Born J. The memory function of sleep. Nat Rev Neurosci. 2010;11:114–126.

25. Diesmann M, Gewaltig MO, Aertsen A. Stable propagation of synchronous spiking in cortical neural networks. Nature. 1999;402:529–533.

26. Dragoi G, Tonegawa S. Distinct preplay of multiple novel spatial experiences in the rat. Proc Natl Acad Sci USA. 2013;110:9100–9105.

27. English DF, Peyrache A, Stark E, Roux L, Vallentin D, Long MA, Buzsàki G. Excitation and inhibition compete to control spiking during hippocampal ripples: intracellular study in behaving mice. J Neurosci. 2014;34:16509–16517.

28. Felsen G, Touryan J, Han F, Dan Y. Cortical sensitivity to visual features in natural scenes. PLoS Biol. 2005;3:e342.

29. Feng T, Silva D, Foster DJ. Dissociation between the experience-dependent development of hippocampal theta sequences and single-trial phase precession. J Neurosci. 2015;35:4980–4902.

30. Foster DJ, Wilson MA. Reverse replay of behavioural sequences in hippocampal place cells during the awake state. Nature. 2006;440:680–683.

31. Gerstner W. Time structure of the activity in neural network models. Phys Rev E. 1995;51:738–758.

32. Gerstner W, Kempter R, van Hemmen JL, Wagner H. A neuronal learning rule for sub-millisecond temporal coding. Nature. 1996;386:76–78.

33. Gerstner W, Kistler WM. Spiking neuron models: Single neurons, population, plasticity. Cambridge, UK: Cambridge UP.;2002.

34. Goodman DFM, Brette R. The brian simulator. Front Neurosci. 2009;3:192–197.

35. Graupner M, Brunel N. Calcium-based plasticity model explains sensitivity of synaptic changes to spike pattern, rate, and dendritic location. Proc Natl Acad Sci USA. 2012;109:3991–3996.

36. Halff AW, Géomez-Varela D, John D, Berg DK. A novel mechanism for nicotinic potentiation of glutamatergic synapses. J Neurosci. 2014;34:2051–2064.

37. Hahn G, Bujan AF, Fréegnac Y, Aertsen A, Kumar A. Communication through resonance in spiking neuronal networks. PLoS Comput Biol. 2014;10:e1003811.

38. Hanse E, Seth H, Riebe I. AMPA-silent synapses in brain development and pathology. Nat Rev Neurosci. 2013;14:839–850.

39. Hasselmo ME. Neuromodulation: acetylcholine and memory consolidation. Trends Cogn Sci. 1999;3:351–359.

40. Hasselmo ME, Schnell E, Barkai E. Dynamics of learning and recall at excitatory recurrent synapses and cholinergic modulation in rat hippocampal region CA3. J Neurosci. 1995;15:5249–5262.

41. Hebb DO. The organization of behavior: A neuropsychological theory. New York.;1949.

42. Hennequin G, Vogels TP, Gerstner W. Non-normal amplification in random balanced neuronal networks. Phys Rev E. 2012;86:011909.

43. Horn D, Levy N, Meilijson I, Ruppin E. Distributed synchrony of spiking neurons in a Hebbian cell assembly. Adv Neural Inf Process Syst. 2000;12:129–135.

44. Jahnke S, Memmesheimer RM, Timme M. Propagating synchrony in feed-forward networks. Front Comput Neurosci. 2013;7:153.

45. Kammerer A, Tejero-Cantero A´, Leibold C. Inhibition enhances memory capacity: optimal feedback, transient replay and oscillations. J Comput Neurosci. 2013;34:125–136.

46. Kappel D, Nessler B, Maass W. STDP installs in winner-take-all circuits an online approximation to hidden Markov model learning. PLoS Comput Biol. 2014;10:e1003511.

47. Karlsson MP, Frank LM. Awake replay of remote experiences in the hippocampus. Nature Neurosci. 2009;12:913–918.

48. Kempter R, Gerstner W, van Hemmen JL. Hebbian learning and spiking neurons. Phys Rev E. 1999;59:4498.

49. Kenet T, Bibitchkov D, Tsodyks M, Grinvald A, Arieli A. Spontaneously emerging cortical representations of visual attributes. Nature. 2003;425:954–956.

50. Klampfl S, Maass W. Emergence of dynamic memory traces in cortical microcircuit models through STDP. J Neurosci. 2013;33:11515–11529.

51. Ko H, Hofer SB, Pichler B, Buchanan KA, Sjäosträom PJ, Mrsic-Flogel TD. Functional specificity of local synaptic connections in neocortical networks. Nature. 2011;473:87–91.

52. Kowalski J, Gan J, Jonas P, Pernéıa-Andrade AJ. Intrinsic membrane properties determine hippocampal differential firing pattern in vivo in anesthetized rats. Hippocampus. 2015;26:668–682.

53. Kruskal PB, Li L, MacLean JN. Circuit reactivation dynamically regulates synaptic plasticity in neocortex. Nat Commun. 2013;4:2574.

54. Kumar A, Rotter S, Aertsen A. Conditions for propagating synchronous spiking and asynchronous firing rates in a cortical network model. J Neurosci. 2008;28:5268–5280.

55. Kumar A, Rotter S, Aertsen A. Spiking activity propagation in neuronal networks: reconciling different perspectives on neural coding. Nature Rev Neurosci. 2010;11:615–627.

56. Lashley KS. The problem of serial order in behavior. In: Cerebral mechanisms in behavior (Jeffress LA, ed), pp 112–131. New York: Wiley.;1951.

57. Lazar A, Pipa G, Triesch J. SORN: a self-organizing recurrent neural network. Front Comput Neurosci. 2009;3:23.

58. Lee AK, Wilson MA. Memory of sequential experience in the hippocampus during slow wave sleep. Neuron. 2002;36:1183–1194.

59. Leibold C, Kempter R. Memory capacity for sequences in a recurrent network with biological constraints. Neural Comput. 2006;18:904–941.

60. Leibold C, Kempter R. Sparseness constrains the prolongation of memory lifetime via synaptic metaplasticity. Cereb Cortex. 2008;18:67–77.

61. Litwin-Kumar A, Doiron B. Slow dynamics and high variability in balanced cortical networks with clustered connections. Nat Neurosci. 2012;15:1498–1505.

62. Litwin-Kumar A, Doiron B. Formation and maintenance of neuronal assemblies through synaptic plasticity. Nat Commun. 2014;5:5319.

63. Luczak A, Bartho` P, Harris KD. Spontaneous events outline the realm of possible sensory responses in neocortical populations. Neuron. 2009;62:413–425.

64. Malenka RC, Bear MF. LTP and LTD: an embarrassment of riches. Neuron. 2004;44:5–21.

65. Marrosu F, Portas C, Mascia MS, Casu MA, F`a M, Giagheddu M, Imperato A, Gessa GL. Microdialysis measurement of cortical and hippocampal acetylcholine release during sleep-wake cycle in freely moving cats. Brain Res. 1995;671:329–332.

66. Mehring C, Hehl U, Kubo M, Diesmann M, Aertsen A. Activity dynamics and propagation of synchronous spiking in locally connected random networks. Biol Cybern. 2003;88:395–408.

67. Memmesheimer RM. Quantitative prediction of intermittent high-frequency oscillations in neural networks with supralinear dendritic interactions. Proc Natl Acad Sci USA. 2010;107:11092–11097.

68. Mishra RK, Kim S, Guzman SJ, Jonas P. Symmetric spike timing-dependent plasticity at CA3-CA3 synapses optimizes storage and recall in autoassociative networks. Nat Commun. 2016;7:11552.

69. Moldakarimov S, Bazhenov M, Sejnowski TJ. Feedback stabilizes propagation of synchronous spiking in cortical neural networks. Proc Natl Acad Sci USA. 2015;112:2545–2550.

70. Morrison A, Aertsen A, Diesmann M. Spike-timing-dependent plasticity in balanced random networks. Neural Comput. 2007;19:1437–1467.

71. Mostafa H, Indiveri G. Sequential activity in asymmetrically coupled winner-take-all circuits. Neural Comput. 2014;26:1973–2004.

72. Murphy BK, Miller KD. Balanced amplification: a new mechanism of selective amplification of neural activity patterns. Neuron. 2009;61:635–648.

73. Okun M, Lampl I. Instantaneous correlation of excitation and inhibition during ongoing and sensory-evoked activities. Nat Neurosci. 2008;11:535–537.

74. Omura Y, Carvalho MM, Inokuchi K, Fukai T. A lognormal recurrent network model for burst generation during hippocampal sharp waves. J Neurosci. 2015;35:14585–145601.

75. Pfister JP, Gerstner W. Triplets of spikes in a model of spike timing-dependent plasticity. J Neurosci. 2006;26:9673–9682.

76. Rapp PR, Gallagher M. Preserved neuron number in the hippocampus of aged rats with spatial learning deficits. Proc Natl Acad Sci USA. 1996;93:9926–9930.

77. Renart A, Moreno-Bote R, Wang XJ, Parga N. Mean-driven and fluctuation-driven persistent activity in recurrent networks. Neural Comput. 2007;19:1–46.

78. Rezende DJ, Gerstner W. Stochastic variational learning in recurrent spiking networks. Front Comput Neurosci. 2014;8:38.

79. Ricciardi LM. Diffusion processes and related topics on biology. Berlin: Springer.;1977.

80. Romani S, Tsodyks M. Short-term plasticity based network model of place cells dynamics. Hippocampus. 2015;25:94–105.

81. Roudi Y, Latham PE. A balanced memory network. PLoS Comput Biol. 2007;3:1679–1700.

82. Sadeh S, Clopath C, Rotter S. Emergence of functional specificity in balanced networks with synaptic plasticity. PLoS Comput Biol. 2015;11:e1004307.

83. Sadovsky AJ, MacLean JN. Scaling of topologically similar functional modules defines mouse primary auditory and somatosensory microcircuitry. J Neurosci. 2013;33:14048–14060.

84. Scarpetta S, de Candia A. Alternation of up and down states at a dynamical phase-transition of a neural network with spatiotemporal attractors. Front Syst Neurosci. 2014;8:88.

85. Song S, Sjäosträom PJ, Reigl M, Nelson S, Chklovskii DB. Highly nonrandom features of synaptic connectivity in local cortical circuits. PLoS Biol. 2005;3:e68.

86. Stark E, Roux L, Eichler R, Buzséaki G. Local generation of multineuronal spike sequences in the hippocampal CA1 region. Proc Natl Acad Sci USA. 2005;112:10521–10526.

87. Thomas SA. Neuromodulatory signaling in hippocampus-dependent memory retrieval. Hippocampus. 2015;25:415–431.

88. Titchener EB. Lectures on the experimental psychology of the thought-processes. New York: Macmillan.;1909.

89. Trengove C, vanLeeuwen C, Diesmann M. High-capacity embedding of synfire chains in a cortical network model. J Comput Neurosci. 2013;34:185–209.

90. Treves A, Rolls ET. What determines the capacity of autoassociative memories in the brain? Network: Comput Neural Syst. 1991;2:371–397.

91. Tsodyks MV, Markram H. The neural code between neocortical pyramidal neurons depends on neurotransmitter release probability. Proc Natl Acad Sci USA. 1997;94:719–723.

92. van Vreeswijk C, Sompolinsky H. Chaos in neuronal networks with balanced excitatory and inhibitory activity. Science. 1996;274:1724–1726.

93. van Vreeswijk C, Sompolinsky H. Chaotic balanced state in a model of cortical circuits. Neural Comput. 1998;10:1321–1371.

94. Vogels TP, Sprekeler H, Zenke F, Clopath C, Gerstner W. Inhibitory plasticity balances excitation and inhibition in sensory pathways and memory networks. Science. 2011;334:1569–1573.

95. Waddington A, Appleby PA, De Kamps M, Cohen N. Triphasic spike-timing-dependent plasticity organizes networks to produce robust sequences of neural activity. Front Comput Neurosci. 2012;6:88.

96. Wallace DJ, Kerr JND. Chasing the cell assembly. Curr Opin Neurobiol. 2010;20:296–305.

97. Washburn MF. Movement and mental imagery: outlines of a motor theory of the complexer mental processes. Boston: Houghton Mifflin.;1916.

98. West MJ, Slomianka LHJG, Gundersen HJG. Unbiased stereological estimation of the total number of neurons in the subdivisions of the rat hippocampus using the optical fractionator. Anat Record. 1991; 231:482–497.

99. Willshaw DJ, Buneman OP,Longuet-Higgins HC. Non-holographic associative memory. Nature. 1969;222:960–962.

100. Wilson MA, McNaughton BL. Dynamics of the hippocampal ensemble code for space. Science. 1993;261:1055–1058.

101. Xue M, Atallah BV, Scanziani M. Equalizing excitation-inhibition ratios across visual cortical neurons. Nature. 2014;511:596–600.

102. Zenke F, Agnes EJ, Gerstner W. Diverse synaptic plasticity mechanisms orchestrated to form and retrieve memories in spiking neural networks. Nature Commun. 2015;6:6922.

